# Decoding universal principles of codon-mediated regulation of gene expression

**DOI:** 10.64898/2026.05.27.728126

**Authors:** Daisuke Tsugama, Kota Kambara

## Abstract

Codon usage determines gene expression levels, yet its universal principles remain elusive. Here, we developed a regression-based model to derive “codon weights” from transcriptomic data, enabling improved prediction of mRNA and protein abundance across diverse taxa, including plants, mammals, insects, and microbes. Ribosome profiling (Ribo-seq) data analysis revealed that these codon weights correlate with Ribo-seq-weighted cumulative codon frequencies specifically in ribosome-unoccupied regions, rather than at stalling sites, across all seven model species. Experimental validation using species-specific optimization confirmed that our method effectively modulates gene expression in *Escherichia coli* and terrestrial plants. These findings demonstrate that species-specific environments for gene expression are encoded in codon weights, which can be deduced through a universal, species-independent framework, providing a new foundation for synthetic biology.

## Introduction

Codon usage is a determinant of translation efficiency, with its impact varying across both specific codons and biological species. These effects are thought to be governed by the abundance and specificity of diverse regulatory factors, including ribosome translocation modulators (e.g., tRNAs and RNA helicases) (1, 2) and mRNA stability regulators (e.g., RNases and RNA-binding proteins) (2–6). For biotechnological and research applications, codon optimization of a coding sequence (CDS) is a widely used strategy to enhance protein expression in specific hosts (7, 8). Several codon-related indices have been developed to evaluate these effects, such as the GC content at the third position (GC3) (9, 10), the codon adaptation index (CAI) (8, 11, 12), the tRNA adaptation index (tAI) (13) and its derivatives, stAI (species-specific tAI) (14) and gtAI (genetic tAI) (15). Although these indices correlate with protein abundance in specific organisms, their performance has not been systematically compared across a broad species, organs, or developmental stages. Consequently, establishing an effective and universal framework for codon-related indices, particularly for plants, where such evaluations are limited, remains a challenge.

To address this gap, RNA sequencing (RNA-seq)-derived transcriptome data were analyzed to obtain mRNA abundance for a variety of samples primarily for *Arabidopsis thaliana*, rice (*Oryza sativa*), humans (*Homo sapiens*) and mice (*Mus musculus*), plant and animal model species that were most rich in such data. Protein abundance data were also collected for various samples from these species. While applying the existing codon-related indices and trying to generate new indices, codon frequencies were found to correlate with mRNA and protein abundance in numerous samples across all of the four species. These observations prompted the development of a model based on the regression of mRNA abundance (transcripts per million; TPM) against codon frequencies to obtain predicted TPM as a novel codon-related index. In this model, the regression coefficients assigned to each codon were defined as “codon weights”, representing the relative contribution of each codon frequency to the predicted mRNA abundance. Extended acquisition and characterization of codon weights, integrated with ribosome profiling (Ribo-seq, (16, 17)) data, reveal a universal principle regarding their sites of action and demonstrate their broad, species-independent applicability to predicting and modulating gene expression.

## Results

### Codon weights derived from simple regression effectively predict gene expression levels

Five regression models (simple linear, Ridge, Lasso, weighted, and principal component) were evaluated in four representative plant and animal species (*Arabidopsis*, rice, humans, and mice). The resulting predicted TPM values exhibited robust correlations with observed TPM, mRNA half-lives, and protein abundance across most samples in all species, with no substantial performance differences observed between the models (fig. S1). The simple linear regression model was employed for subsequent analyses due to its simplicity and interpretability. Consistent with the model design, the predicted TPM values exhibited stronger statistical associations with observed TPM, mRNA half-life, and protein abundance than individual codon frequencies. These values also generally outperformed existing codon indices (Fig. 1A and fig. S2). The predicted TPM values for three additional model species (fruit fly, budding yeast and *Escherichia coli*) also correlated with observed protein abundance (fig. S3). In a comparison across all seven studied species, the predicted TPM values outperformed GC3 content in all cases, showed higher performance than the CAIhigh (CAI based on highly expressed genes) in *Arabidopsis*, rice, humans, and mice while remaining comparable to the CAIhigh in the other three species, and showed performance second only to the stAI in humans (Fig. 1B).

**Figure 1.**
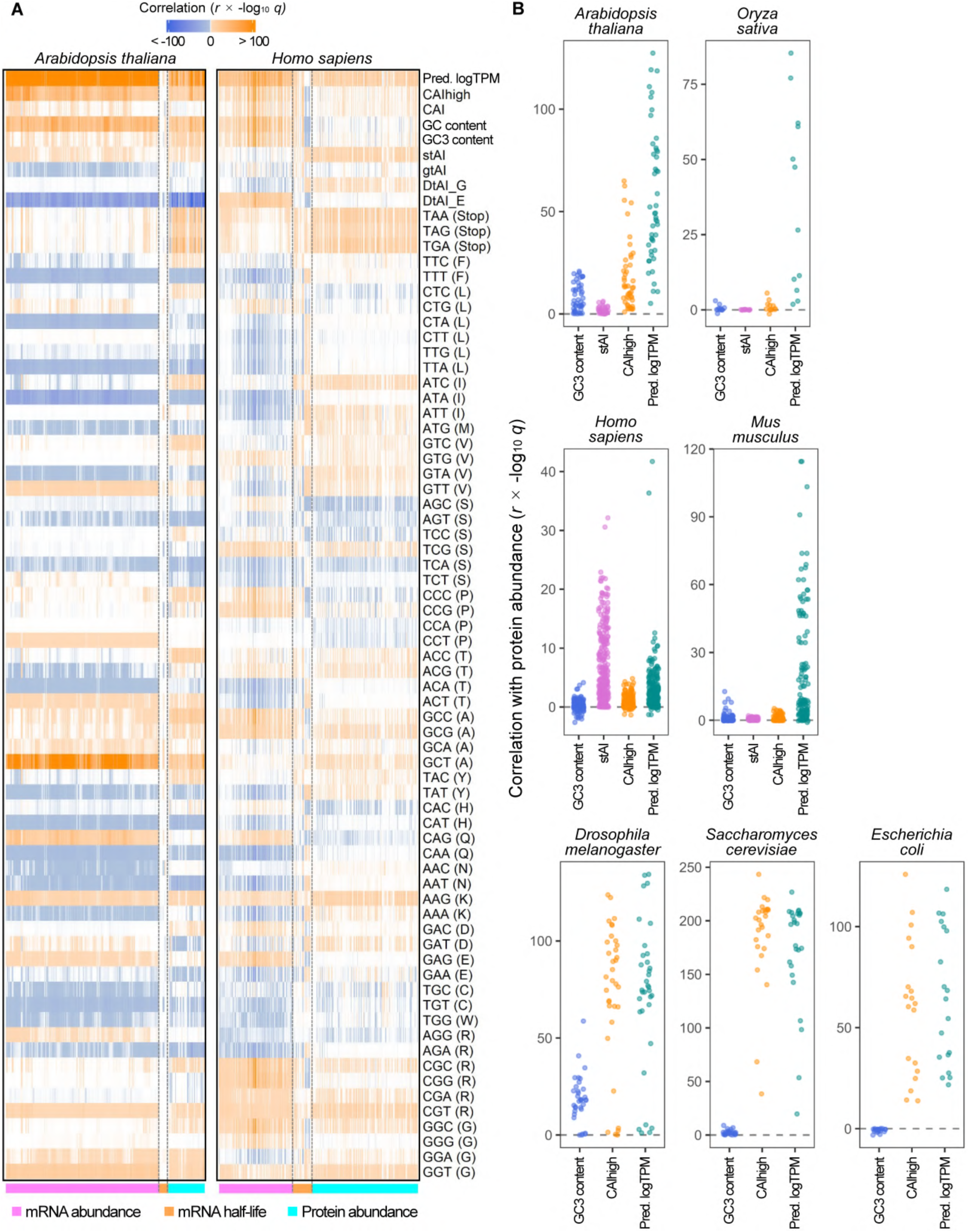
Codon frequencies are determinants of mRNA and protein abundance. (A) Heatmaps of Pearson’s *r* multiplied by -log10 *q* (FDR-corrected *P* values) calculated from correlations of predicted log10(TPM) (Pred. logTPM), codon-related indices, and individual codon frequencies with observed log10(TPM) (mRNA abundance), mRNA half-life, and log10(protein abundance). Predicted values were obtained from simple linear regression models trained on genes with TPM > 0. (B) Jitter plots of Pearson’s *r* multiplied by -log10 *q* (FDR-corrected *P* values) calculated from correlations of GC3 content, stAI (only for *A. thaliana*, *O. sativa*, *H. sapiens*, and *M. musculus*), CAIhigh, and Pred. logTPM with observed log10(protein abundance). Observed log10(protein abundance) was used as the reference for all correlations. Species names are indicated at the top of the panels in both (A) and (B).

The analysis was expanded to 29 species, primarily comprising terrestrial plants and plant-associated microorganisms, to further evaluate the applicability of the regression-based model. The predicted TPM values exhibited strong statistical associations with observed TPM across all 29 species (fig. S4) and outperformed GC3 content, CAIhigh, and stAI in all tested species (fig. S5).

UMAP visualization revealed that taxonomically related species have similar codon weight profiles and that dicotyledonous plants, including *Arabidopsis*, form a distinct cluster (fig. S6). Hierarchical clustering identified specific codons, including ATG, AAG, and those consisting only of A and T, as contributors to the taxonomic differences in codon weight profiles (Fig. 2). These results suggest that codon weights, which are tailored to individual species and taxon, can be captured by the universally applicable regression-based model.

**Figure 2.**
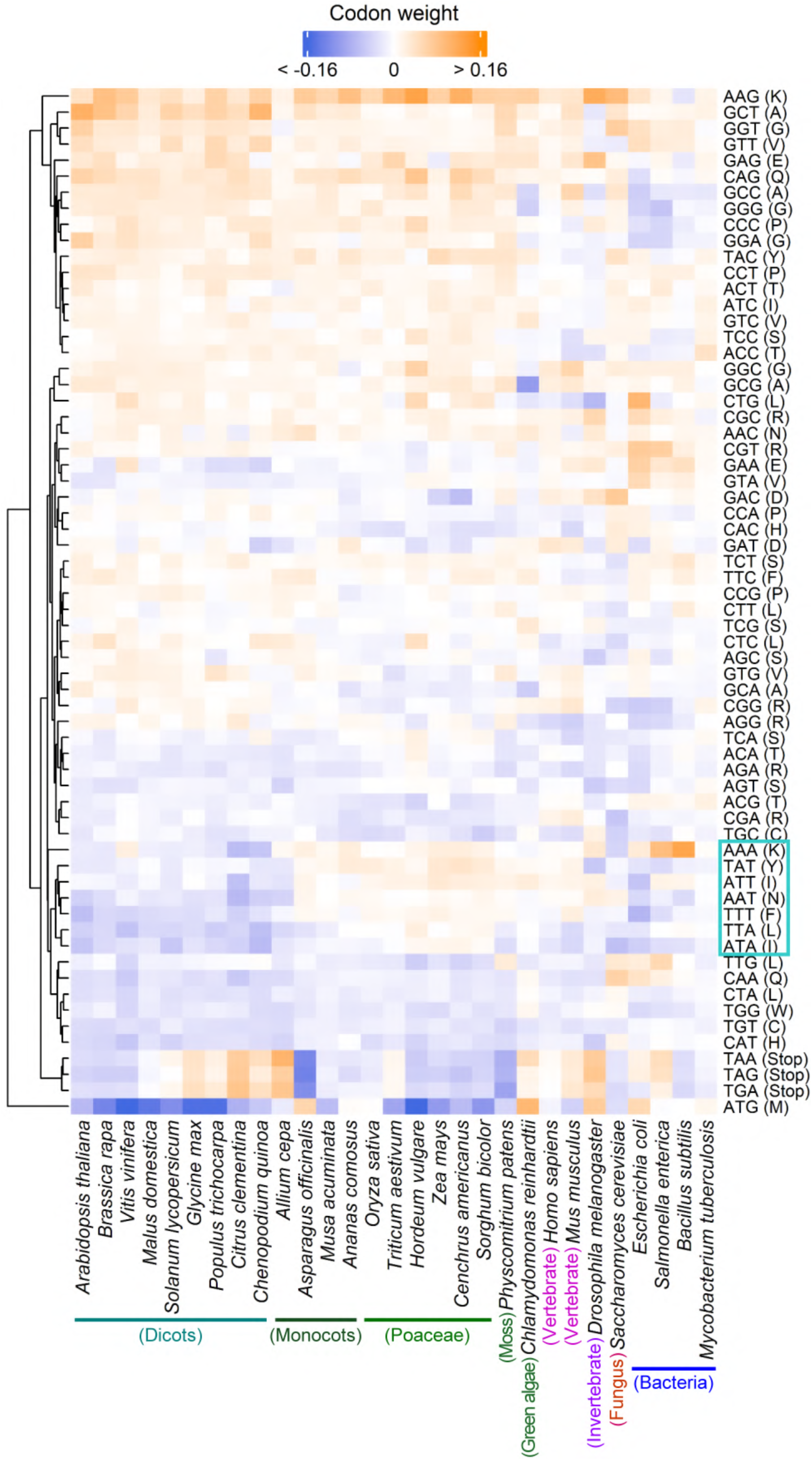
Codon weights from regression of mRNA abundance across 29 species. The codon weights were obtained as coefficients from simple linear regression models trained on genes with TPM > 0. Codons were clustered based on their weight profiles. Species were ordered primarily by phylogenetic classification and secondarily by the similarity of their codon weights. Seven codons consisting solely of A and T are boxed on the right. All values are presented without scaling.

### Codon weights correlate with codon frequencies specifically in ribosome-unoccupied regions

To investigate the mechanistic basis of the codon weight-based predictions, Ribo-seq data were analyzed, obtaining the Ribo-seq-weighted cumulative codon frequencies (hereafter referred to as weighted frequencies) at 101 sites centered on the mapped P-site. Heatmaps revealed mosaic patterns of weighted codon frequencies only at the putative P-site and its proximal regions (approximately ± 5 sites), reflecting the physical occupancy of the ribosome across all seven species studied (Fig. 3A and fig. S7). In the four eukaryotic species (plants and animals), weighted stop codon frequencies were specifically enriched at the A-site compared to other sites, consistent with ribosome stalling during translation termination (16); this trend was less prominent in the other three species (fig. S7). Analysis of codon groups based on GC content or the third letter revealed codon group-specific, ribosome occupancy-dependent, and species-specific peaks across the 101 sites. These peaks appeared relevant to the ribosome entry, A-, P-, E-, and exit sites, and became more prominent and acute, with their positions unchanged, when the analysis was restricted to sites with stronger ribosome stalling (Fig. 3B, fig. S8, and fig. S9).

**Figure 3.**
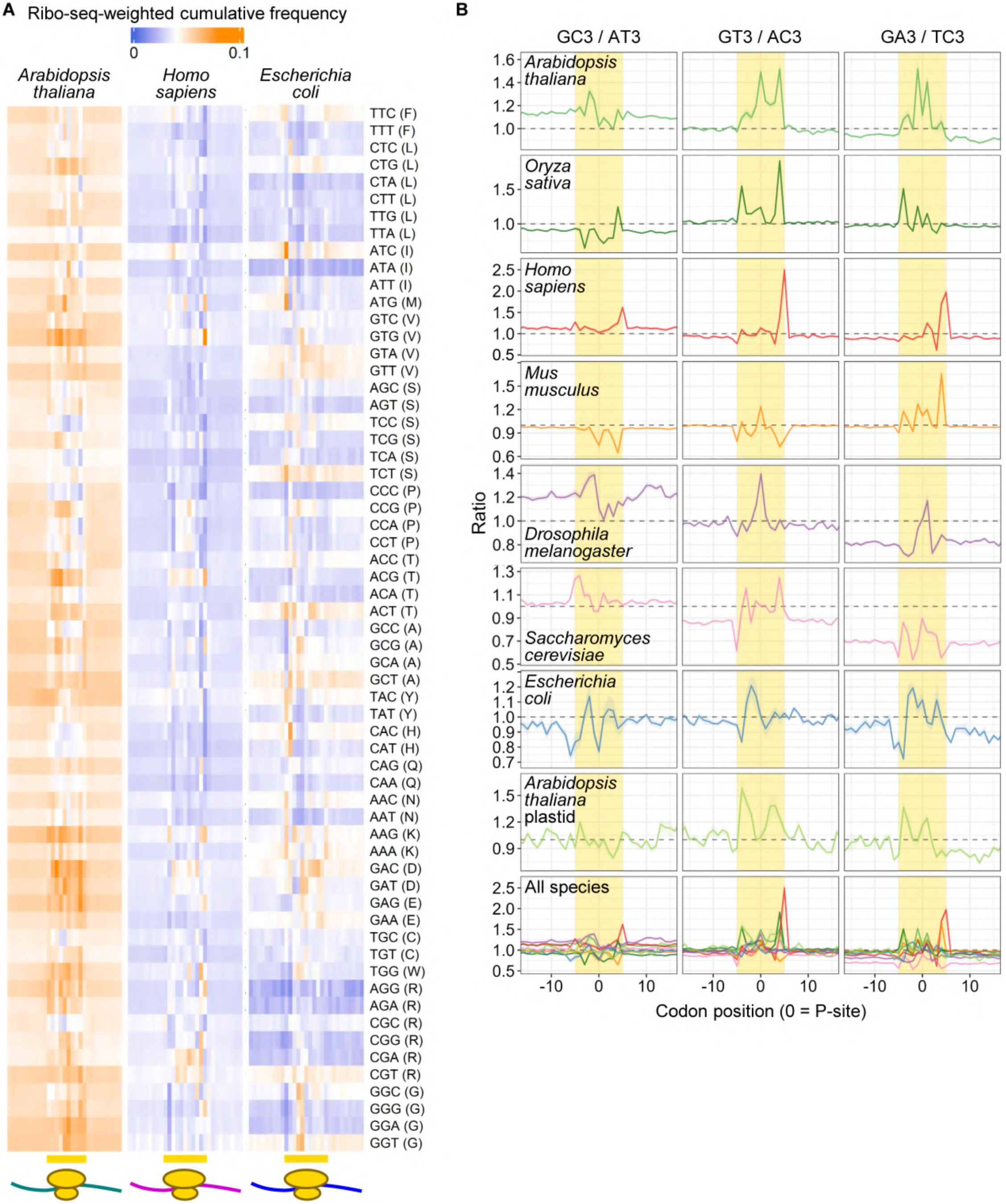
Ribo-seq-weighted codon frequencies differ between ribosome-occupied and unoccupied sites on mRNA across species. (A) Heatmap of Ribo-seq-weighted cumulative codon frequencies for *Arabidopsis thaliana*, *Homo sapiens*, and *Escherichia coli*. At each position within a 101-codon window centered on reference sites (offsets), P-site read counts were normalized by the mean read density of each gene, summed up for each codon across all genes, and subsequently divided by the total occurrence of each codon across all analyzed genes. Values for the putative P-site (site 51, defined in fig. S7) and its 14 flanking upstream and downstream sites are presented. Putative ribosome-occupied sites are indicated by bars and ribosome icons at the bottom. (B) Site-specific nucleotide composition ratios. Ratios were calculated for synonymous codon groups categorized by their third-position bases (e.g., GC3/AT3 represents the ratio of codons ending in G or C to those ending in A or T). Values were calculated for 29 sites centered on the P-site using the cumulative frequencies described in (A). Data for the stalling+ subsets (defined in fig. S9) across seven species are presented as ribbon plots (mean ± range [min–max]). The minimal variance observed highlights the high conservation of codon composition patterns around stalling sites.

In all seven species except rice, codon weights exhibited strong statistical associations with weighted codon frequencies only in ribosome-unoccupied regions. In rice, such associations were observed specifically when the analysis was restricted to AT3 codons. The AT3/GC3 codon grouping appeared relevant to the codon weight-frequency associations in *Arabidopsis*, humans, and mice (Fig. 4A), whereas the GA3/TC3 grouping was more prominent in fruit fly and *E. coli* (fig. S10). Representative scatter plots confirmed the site-specific bias of codon group ratios and the ribosome-unoccupied region-specific associations between codon weights and weighted frequencies (Fig. 4B). These results indicate that species-specific factors define ribosome-occupied (or ribosome-stalling) regions and that codon-dependent regulation of gene expression primarily functions outside these regions across species.

**Figure 4.**
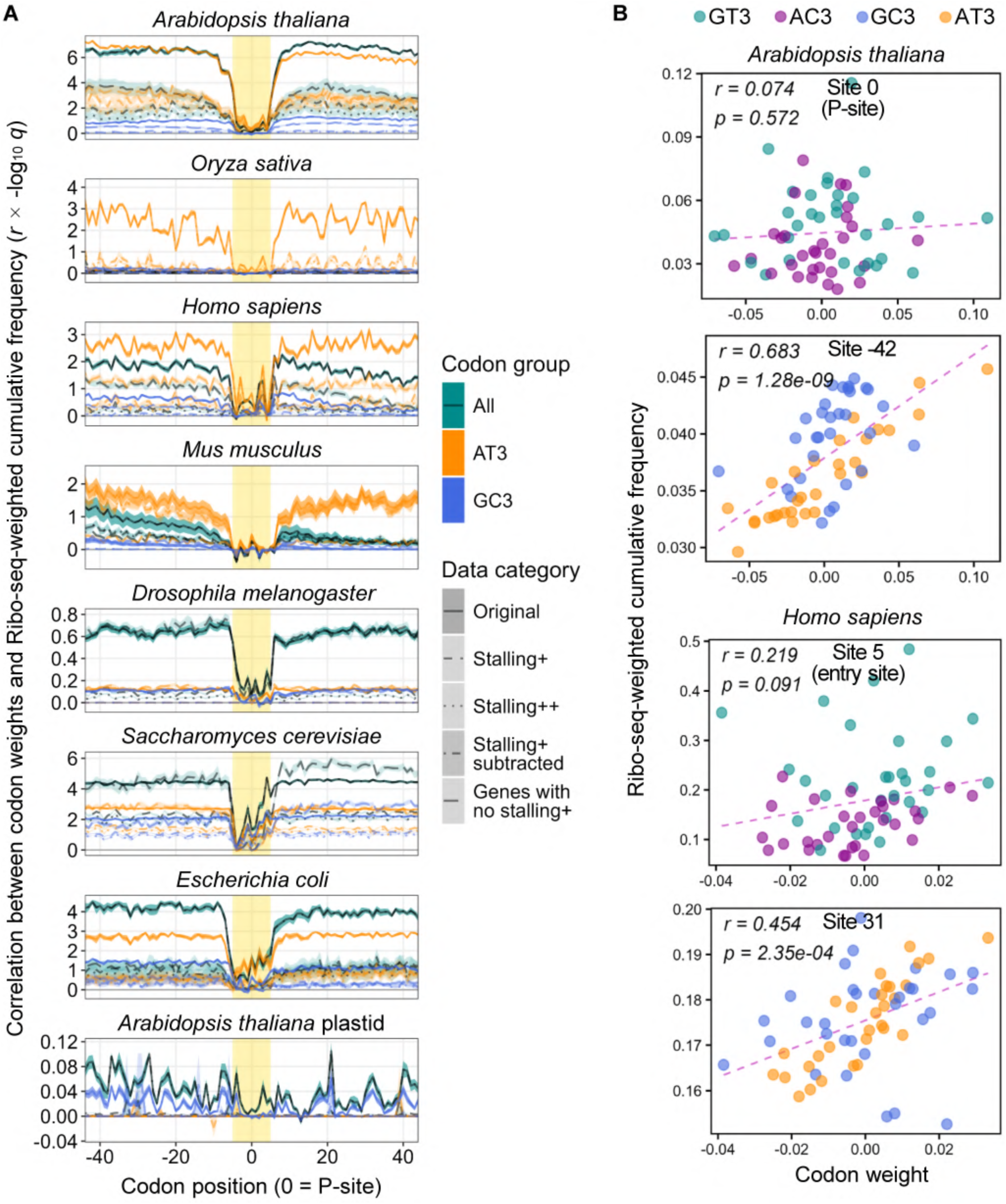
Codon weights correlate with Ribo-seq-weighted codon frequencies at ribosome-unoccupied sites across species. (A) Positional correlations between codon weights and Ribo-seq-weighted cumulative codon frequencies. Values of Pearson’s *r* multiplied by -log10 *q* (FDR-corrected *P* values) were calculated by correlating the codon weights (from simple linear regression models trained on genes with TPM > 0) with the normalized codon counts across an 81-codon window. For each position, the correlation was tested using the 61 non-stop codons. These values were determined independently for each Ribo-seq sample and are presented as ribbon plots representing the range between the minimum and maximum values, with the middle line indicating the mean. The putative ribosome-occupied regions were estimated by the codon frequency patterns and the putative P-sites presented in fig. S7 and are highlighted by background color. Species names are indicated at the top of the panels. Similar plots for different codon groups are provided in fig. S10. (B) Scatter plots of codon weights versus Ribo-seq-weighted cumulative codon frequencies at representative sites in *Arabidopsis thaliana* (top panel) and *Homo sapiens* (bottom). The putative P-site in *A. thaliana* (site 0) and the putative mRNA entry site in *H. sapiens* (site 5) were selected as sites that exhibited a high GT3/AC3 ratio (cf. fig. 3B) and no weight-frequency correlation. Site −42 for *A. thaliana* and site 31 for *H. sapiens* were selected as reference sites where the weight-frequency correlation is most significant. Pearson’s correlation coefficient (*r*) and P values (*p*) are indicated within each panel.

### Implementation of codon weight-based optimization for modulating gene expression

To test the practical utility of the codon weight-based prediction method, species-specific codon optimization was applied to two marker genes: *HPT-GFP* and *picALuc2.0-E50K* (hereafter “*picALuc*”). *HPT-GFP* variants were designed to simultaneously target two species using a maximin strategy, whereas *picALuc* variants were designed for single species (Fig. 5A). In *Escherichia coli*, GFP signals and hygromycin-resistant colonies were detected only from the yeast/*E. coli*-optimized *HPT-GFP* variant, whereas no signal was observed from either the suboptimal or *Arabidopsis*/onion-optimized variant (Fig. 5B, left panels). Similarly, the highest picALuc signal intensity was obtained from the yeast/*E. coli*-optimized variant (Fig. 5B, right). In contrast, in *Saccharomyces cerevisiae* (budding yeast), hygromycin-resistant colonies were obtained from the suboptimal *HPT-GFP* variant, but not from the yeast/*E. coli*-optimized version. No significant differences in picALuc signal intensities were observed among the codon variants in yeast, either (fig. S11). In onion, GFP signal intensities were comparable among the *HPT-GFP* variants, but were highest for the *Arabidopsis*/onion-optimized *GFP* variant (lacking *HPT*) (Fig. 5C). Similarly, in bok choy (*Brassica rapa* subsp. *chinensis*), which is taxonomically related to *Arabidopsis*, the *Arabidopsis*/onion-optimized *GFP* variant yielded the highest signal intensity, while the suboptimal variant resulted in the lowest mRNA level (Fig. 5D, left). In contrast, no significant differences were observed among the *picALuc* variants in either signal intensities or mRNA levels in bok choy (Fig. 5D, right). The impact of codon optimization was further assessed using in vitro transcription-translation systems, where certain cellular regulators of mRNA and protein abundance are absent. In the wheat germ extract system, no significant differences were observed in picALuc signal intensities among the variants (Fig. 5E, left). In the rabbit reticulocyte lysate system, picALuc signal intensities were lowest for the wheat-optimized variant and comparable among the others (Fig. 5E, right); these results were unaffected by the presence of RNase inhibitors (Fig. 5E). The effect of protein fusion was also examined using *HPT-GFP* variants fused to a common glutathione S-transferase (GST) sequence (*GST-HPT-GFP*). These fusions exhibited attenuated differences in predicted TPM values compared to the unfused versions, but GFP signals in *E. coli* were detected only from the yeast/*E. coli*-optimized variant (fig. S12). However, hygromycin-resistant colonies were obtained from both the yeast/*E. coli*-optimized and the suboptimal variants (fig. S12). These results indicate that, although the effects of codon weight-based optimization depend on specific gene and host contexts, such optimization can effectively improve mRNA and/or protein abundance, particularly in *E. coli*.

**Figure 5.**
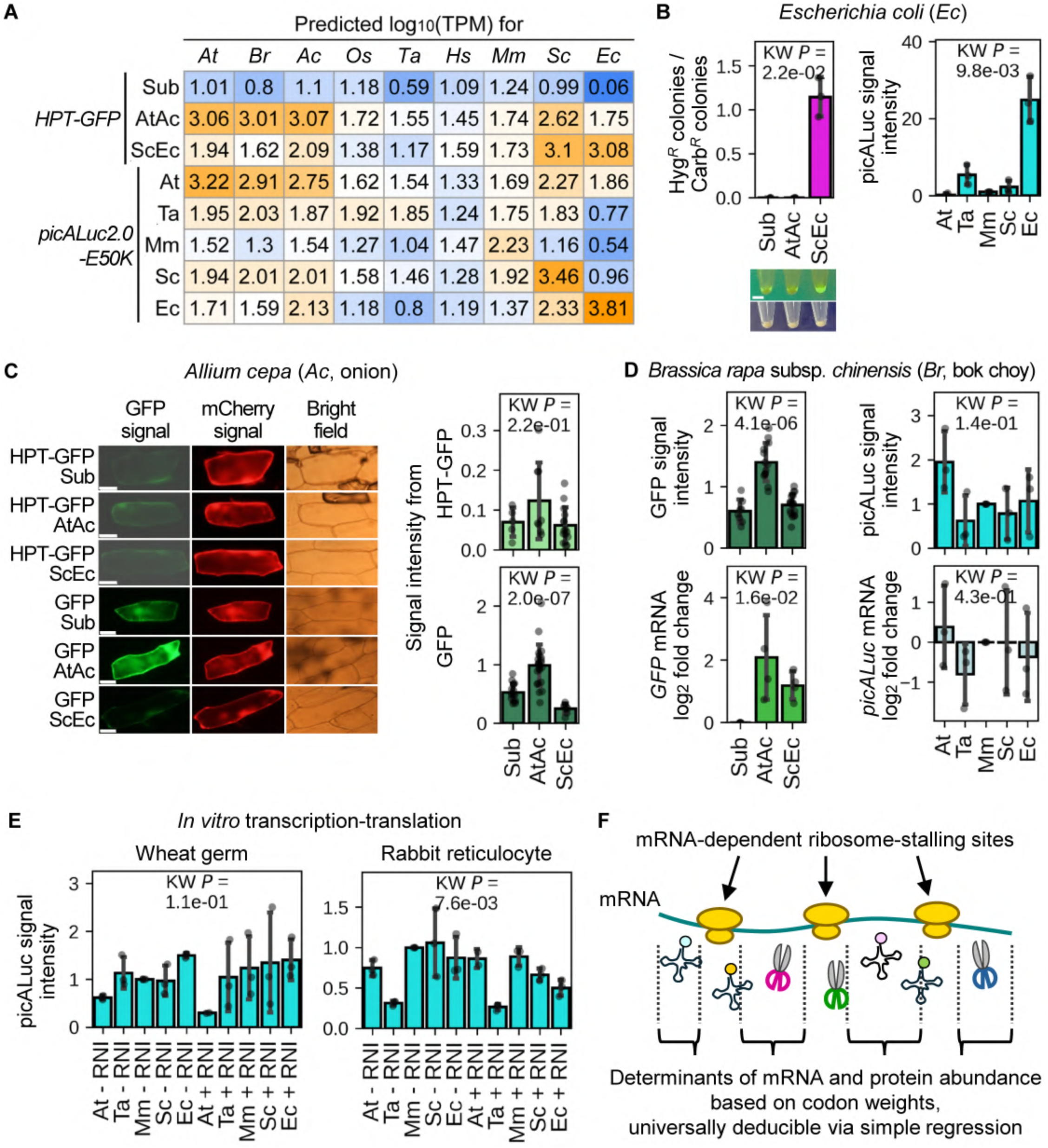
Effects of codon weight-based codon optimization on gene expression. (A) Predicted log10(TPM) of codon variants of *HPT-GFP* and *picALuc2.0-E50K* (hereafter ‘picALuc’). *HPT-GFP* variants include Sub (suboptimal), AtAc (optimized for *A. thaliana* and *Allium cepa*), and ScEc (for *S. cerevisiae* and *E. coli*). *picALuc* variants include At (optimized for *A. thaliana*), Ta (for *T. aestivum*), Mm (for *M. musculus*), Sc (for *S. cerevisiae*) and Ec (for *E. coli*). TPM values were predicted using simple linear regression models trained on genes with TPM > 0 for *A. thaliana* (*At*), *B. rapa* (*Br*), *Allium cepa* (*Ac*), *O. sativa* (*Os*), *T. aestivum* (*Ta*), *H. sapiens* (*Hs*), *M. musculus* (*Mm*), *S. cerevisiae* (*Sc*) and *E. coli* (*Ec*). (B) Performance of codon variants in *E. coli*. *HPT-GFP* and *picALuc* variants were transformed into *E. coli* strain BL21(DE3). Left: Ratios of hygromycin-resistant (Hyg*^R^*; indicating *HPT* functionality) to carbenicillin-resistant (Carb*^R^*; control) colonies, followed by representative GFP signal intensities from cultured cells. Scale bar = 4 mm. Right: picALuc signal intensities in cultured cells. (C) Performance of codon variants in onion (*Allium cepa*). *HPT-GFP* and *GFP* (derived from *HPT-GFP*) variants were co-expressed with *mCherry* in onion epidermal cells. Left: Representative GFP, mCherry, and bright-field images. Scale bar, 50 μm. Right: HPT-GFP (top) and GFP (bottom) signal intensities normalized to mCherry signals. (D) Performance of codon variants in bok choy (*Brassica rapa* subsp. *chinensis*). *GFP* variants were co-expressed with either *mCherry* or *FLuc*, and *picALuc* variants were co-expressed with *FLuc* in bok choy leaf epidermal cells. Left: GFP signal intensities normalized to mCherry (top) and *GFP* mRNA log2 fold changes normalized to *FLuc* as an internal control (bottom). Right: picALuc signal intensities normalized to FLuc signal intensities (top) and *picALuc* mRNA log2 fold changes normalized to *FLuc* as an internal control gene (bottom). (E) Performance of picALuc variants in *in vitro* transcription-translation systems. picALuc variants were expressed *in vitro* using wheat germ extracts (left) or rabbit reticulocyte lysates (right) in the presence of RNase inhibitor (+ RNI) or its absence (- RNI). For all quantitative plots in (B–E), bars and error bars represent means and SD, respectively. Individual data points (dots) correspond to single cells for GFP signal intensities, and to independent biological replicates for picALuc signal intensities and mRNA quantification. KW *P* indicates the *P* value determined by the Kruskal-Wallis test. (F) Summary of findings in this study. Ribosomes stalled on mRNA transcripts are shown with aminoacyl-tRNAs and RNases (represented by scissors), which can influence translation efficiency and mRNA stability in a species-specific manner. Codon frequencies within ribosome-unoccupied regions (indicated by dashed lines and brackets) govern both mRNA and protein abundance. These effects are predictable using codon weights derived from simple transcriptome-wide regression independently of species, providing a universal principle for regulation of gene expression.

## Discussion

### Biological and practical significance of species-specific codon weights

The regression-based approach established in this study offers several advantages over conventional codon indices. Most importantly, it allows for the derivation of species-specific codon weights solely from RNA-seq data. For species where such data are difficult to obtain, weights can be inferred from taxonomically related species, leveraging the observation that codon weight profiles are largely conserved within related taxa (Fig. 2 and fig. S6). Furthermore, unlike indices such as CAIhigh that rely on sets of highly expressed genes, the regression model incorporates information from the entire genes including low-expression genes. This comprehensive integration likely contributes to its superior predictive performance. Because the current models are simple and interpretable, they can be effectively combined with other regulatory features such as promoter and terminator strength or mRNA secondary structure (8) to further refine the precision of expression level predictions and codon optimization.

### Spatial decoupling of codon effects and ribosome stalling

A pivotal finding of this work is the spatial decoupling between codon weights and ribosome stalling. The observation that codon weights primarily correlate with weighted codon frequencies in ribosome-unoccupied regions (Fig. 4) suggests that the dominant codon-based determinants of gene expression are located in these regions. While ribosome-occupied regions are likely defined by conserved structural constraints and remain physically shielded throughout the lifespan of mRNA, the unoccupied regions are more accessible to regulatory machinery. In these accessible landscapes, codon composition can dictate mRNA stability (10) or modulate translation initiation/re-initiation rates (8) through regulatory factors such as RNases and tRNA availability in a host-specific manner. The regulatory sensitivity to specific codon groups, such as the apparently prominent role of AT3/GC3-richness in *Arabidopsis*, humans, and mice, or its specific restriction to AT3 codons in rice, can reflect how species-specific evolutionary pressures shape the biochemical accessibility of the mRNA template. These environments are all encoded in codon weights, which can be deduced from the presented universal, species-independent model. This model shifts the focus of codon-mediated regulation from elongation-rate bottlenecks to the biochemical accessibility of the mRNA template (Fig. 5F).

### Practical implementation and future perspectives

The utility of the codon weight-based codon optimization was demonstrated by successful expression modulation of target genes in *E. coli* and certain plant tissues. However, the limited impact observed for *HPT* and *picALuc* in yeast and onion highlights current limitations. Compared to GFP, these enzymes may require complex folding trajectories that constrain protein functionality and/or abundance. It is also possible that mRNA secondary structures, factors not yet explicitly integrated into our linear model, override the benefits of codon-based abundance control (8). Nevertheless, the fact that none of the optimized codon variants resulted in decreased expression compared to their counterparts underscores the robustness of this approach. Our discovery that traditional indices such GC3 and CAIhigh can negatively correlate with mRNA abundance in certain taxa (e.g., Poaceae) warns against the universal adoption of these metrics for optimization.

To further refine the precision of this framework for synthetic genomics, future models should explicitly incorporate mRNA structural features. The comprehensive transcriptome-wide datasets generated in this study can provide a foundation for evaluating how such structural features interact with codon weights to determine net mRNA and protein abundance. Moreover, expanding this analysis to a broader range of protein classes with diverse folding requirements will help elucidate the scope of codon-dependent regulation of gene expression. Integrating codon weights with these context-dependent parameters can enable the establishment of a more sophisticated and predictive platform for the precision design of genetic systems.

## Acknowledgments

We thank Dr. Daniel Voytas for providing the pMKV060 plasmid. The authors acknowledge the use of Gemini 1.5 Flash (Google) for writing assistance in improving the clarity and language flow of the manuscript. All AI-generated suggestions were critically reviewed and edited by the authors, who take full responsibility for the final content.

## Author contributions

**Conceptualization:** D.T., K.K.; **Methodology:** D.T., K.K.; **Investigation:** D.T., K.K.; **Formal Analysis:** D.T., K.K.; **Writing - Original Draft:** D.T.; **Writing - Review & Editing:** D.T., K.K.; **Funding Acquisition:** D.T. and K.K.

## Data and materials availability

This study did not generate new next-generation sequencing (NGS) datasets; all RNA-seq and Ribo-seq data were retrieved from public databases (NCBI SRA/DDBJ), with a full list of accession numbers provided in the Master Accession List on Figshare (DOI: https://doi.org/10.6084/m9.figshare.32414811). All processed datasets, trained regression models, and custom analytical scripts are available via Figshare. Plasmids generated in this study are available from the corresponding author upon request.

## Supplementary Materials

**The PDF file includes:**

Materials and Methods

Figs. S1 to S12

Legends for Tables S1 to S6

References (S1-S41)

## Other Supplementary Material for this manuscript includes the following

Tables S1 to S6

## Supplementary materials

## Materials and Methods

### Data overview and statements of ethics and AI usage

Detailed descriptions of each analysis, including specific objectives, major findings, and the correspondence between input datasets and computational scripts for all figures, are summarized in table S1. Custom codes used for data analysis are listed in table S2. These codes were generated or refined using Google Gemini 1.5 Flash (via the web interface) between November 2025 and May 2026. Each script was manually reviewed, debugged, and validated by the authors to ensure computational accuracy and reproducibility. Pre-analyzed quantitative data for tRNA abundance and mRNA half-lives were curated based on the information provided in table S3. The complete master list of RNA sequencing (RNA-seq) and ribosome profiling (Ribo-seq, (S1, S2)) metadata, as well as processed final and intermediate datasets and all analytical scripts, are available via Figshare (DOI: https://doi.org/10.6084/m9.figshare.32414811). These data were originally collected under institutional approvals and informed consent; thus, no additional ethical clearance was required for this study.

### RNA-seq data analysis

Genome sequences, general feature format 3 (GFF3), and coding sequence (CDS) files were obtained from Ensembl (Release 115, for humans and mice) and Ensembl Genomes (Release 62, for the other species except pearl millet and onion) (S3). For pearl millet and onion, such files were obtained from Figshare (https://doi.org/10.6084/m9.figshare.21261129.v1 (S4) and https://doi.org/10.6084/m9.figshare.28079846.v1 (S5), respectively). Raw RNA-seq reads were retrieved from the National Center for Biotechnology Information (NCBI) Sequence Read Archive (SRA) or DNA Data Bank of Japan (DDBJ). Full metadata for the thousands of SRA accessions analyzed, including their corresponding species and study identifiers, are provided in the Master Accession List on Figshare (DOI: https://doi.org/10.6084/m9.figshare.32414811). Reads were mapped to the respective reference genomes using HISAT2 (S6), and gene-level counts were quantified using featureCounts (S7). Counts were converted to transcripts per million (TPM) using custom Perl scripts (S8), which were used as a measure of mRNA abundance for subsequent analyses.

### Evaluation of codon demand and supply

Codon demand was calculated as previously described (S9). Absolute codon demand was defined as the sum of (codon counts × TPM) for each codon across all genes within a sample. Relative demand was then derived by normalizing absolute demand per codon to the sum of total demand across all codons. For codon supply, tRNA gene copy numbers and expression data (reads per million, RPM) were curated from public datasets (table S3; (S10) for *Arabidopsis* and rice tRNA gene counts and expression; (S11) for human tRNA gene counts; (S12) for human tRNA expression; (S13) for mouse tRNA gene counts and expression). Absolute supply for each anticodon was defined as the sum of tRNA gene counts or RPM, which were further normalized by total tRNA gene counts or total RPM across anticodons, respectively, to calculate relative supply at the anticodon level.

### Calculation of codon-related indices and protein abundance data

Protein abundance (PA) data for *Arabidopsis thaliana*, *Oryza sativa*, *Homo sapiens*, *Mus musculus*, *Drosophila melanogaster*, *Saccharomyces cerevisiae*, and *Escherichia coli* were retrieved from PaxDb (S14).

### Calculation of codon-related indices and protein abundance data (continued)

For each corresponding gene, GC content and GC3 (GC content at the third codon position) were calculated using custom Perl scripts.

For *Arabidopsis*, rice, humans, and mice, the species-specific tRNA adaptation index (stAI) (S15, S16) and genetic tAI (gtAI) (S17) were calculated using custom Python scripts based on the CDS and tRNA gene copy number data described above. The stAI for each gene was defined as the geometric mean of relative codon adaptiveness values (*w_i_*, where *_i_* represents the 61 sense codons), derived from tRNA gene copy numbers and wobble base-pairing efficiencies.

To evaluate the potential impact of codon demand, we also calculated a demand-adjusted tAI based on gene numbers (DtAI_G), defined as the geometric mean of *w_i_* divided by the respective codon frequency. Furthermore, a demand-adjusted tAI based on expression (DtAI_E) was derived by substituting tRNA gene copy numbers with tRNA supply metrics and codon frequencies with codon demand values.

The codon adaptation index (CAI) (S18) for all genes was calculated using the Emboss package (S19). The CAI for highly expressed genes (CAIhigh) was determined by analyzing codon frequencies of genes with total TPM values in the top 5th percentile for each species. For humans, mice, *Arabidopsis*, and rice, CAIhigh values were also independently calculated for each individual RNA-seq sample.

### Estimation of codon weights via regression models

To estimate the contribution of each codon to gene expression, regression analyses was performed using custom Python scripts for all 29 species. The response variable was the log-transformed mRNA abundance, calculated as log1p(TPM). For the explanatory variables, codon frequencies for each gene were transformed using the centered log-ratio (CLR) (S20) to address the constraints of compositional data. Five different regression models were evaluated: simple linear regression (S21), Ridge regression (S22), Lasso regression (S23), weighted linear regression as implemented in scikit-learn (S24), and principal component regression (S25).For the primary model construction, the total TPM summed across all RNA-seq samples was used for each gene. For *H. sapiens*, *M. musculus*, *A. thaliana*, and *O. sativa*, regressions were also performed using TPM values from individual RNA-seq samples to enable subsequent correlation analyses. In all cases, only expressed genes (TPM or total TPM > 0) were included in the regression. The regression coefficients obtained from the simple linear model were defined as the codon weights. UMAP (uniform manifold approximation and projection) (S26) was performed to compare codon weight profiles between species. A heatmap was generated using the raw, non-standardized codon weights to visualize the differences in codon weight profiles across species and codons. These weights visualization was performed using R.

### Correlation analysis between codon indices and gene expression metrics

For the 29 studied species, the statistical associations between observed mRNA abundance [log_10_(TPM); only for genes with TPM > 0] and three indices: GC3 content, stAI, and predicted mRNA abundance [predicted log_10_(TPM)] were examined. Correlations were determined using Pearson’s method (Pearson, 1896) (S27). For the seven model species (*A. thaliana*, *O. sativa*, *H. sapiens*, *M. musculus*, *D. melanogaster*, *S. cerevisiae*, and *E. coli*), the correlations between these indices and PA [log_10_(PA)] were also examined.

### Correlation analysis between codon indices and gene expression metrics (continued)

For *H. sapiens*, *M. musculus*, and *A. thaliana*, mRNA half-life data were retrieved from NCBI Gene Expression Omnibus (GEO) using the accessions listed in table S3 ((S28) for humans and mice; (S29-S31) for *Arabidopsis*). Correlations between mRNA half-lives and the aforementioned codon indices or predicted TPM values were also evaluated.

To account for multiple testing, *P*-values obtained from the correlation tests were adjusted using the false discovery rate (FDR) method (S32) to obtain *q*-values. For visualization, heatmaps and jitter plots were generated using a metric defined as the Pearson correlation coefficient multiplied by -log_10_(*q*). Scatter plots comparing predicted versus observed log_10_(TPM) and predicted log_10_(TPM) versus log_10_(PA) were also constructed. All statistical tests and data visualizations were performed in R.

### Ribosome profiling (Ribo-seq) data analysis

Raw Ribo-seq reads for seven species (see Master Accession List available at Figshare for the complete accession list) were mapped to their respective transcriptomes using STAR (S33), based on GFF3/GTF annotations from Ensembl. To determine the ribosome P-site for each read length, we analyzed the three-nucleotide periodicity and identified length-specific offsets (S2) from the mapping start positions using metaplots generated by RiboCode (S34). For *S. cerevisiae* and *E. coli*, where 5’ untranslated sequence information was absent in the GTF files, an initial offset of zero was used, and the putative P-site was subsequently inferred from the resulting heatmap patterns. For *E. coli*, since clear three-nucleotide periodicity was not detectable by standard metaplot criteria, the optimal read lengths and offsets were manually determined by examining the read distribution around start and stop codons for each dataset. While a broader range of datasets was initially processed, only those passing quality controls, including adequate mapping rates and three-nucleotide periodicity, were included in the final analysis.

The P-site (site 51) and its flanking 50 codons (sites 1-101) were extracted for each gene. For each site within this 101-codon window, a count equal to the number of Ribo-seq reads mapped to site 51 was assigned. These counts were first normalized by mean read density (total reads divided by the number of codons in the CDS) of each gene. These gene-normalized counts were then aggregated across all analyzed genes at each position within the window. To correct for codon usage bias, the aggregated normalized counts were subsequently divided by the total occurrence of each codon within the sequences of the same gene set, yielding a background-corrected observed/expected (O/E) ratio (hereafter referred to as Ribo-seq-weighted cumulative codon frequency).

Using the resulting values, heatmaps were generated in R to visualize stalling patterns. Site-specific codon group ratios (i.e., GC3/AT3, GT3/AC3, and GA3/TC3) were also calculated and visualized using ribbon plots. Finally, for each of the 101 sites, the statistical association between the Ribo-seq-weighted codon frequencies and the corresponding codon weights (see “Estimation of codon weights via regression models”) was evaluated using the correlation tests described above. The resulting Pearson correlation coefficients multiplied by -log_10_(*q*) were visualized using ribbon plots in R.

### Design of codon-optimized gene variants

To modulate gene expression, a Python-based optimization tool that predicts the TPM of any CDS using the regression models and maximizes the predicted value through synonymous codon substitutions was developed. To ensure robust optimization and avoid potential overestimation by any single model, the program was designed to be able to simultaneously integrate multiple regression models. In this study, a maximin strategy (S35) was adopted, where the final objective function for maximization was defined as the minimum predicted TPM value among the selected models.

Using the simple linear regression models described above as input, codons of the *HPT-GFP* fusion gene, consisting of the hygromycin phosphotransferase (*HPT*) from pCAMBIA1300 (Abcam) and *GFP* (NCBI GenBank accession LC741218.1) were modified. As a result, two species-pair-optimized variants, *HPT-GFP_AtAc* (optimized for *A. thaliana* and *A. cepa* (onion)) and *HPT-GFP_ScEc* (optimized for *S. cerevisiae* (budding yeast) and *E. coli*), were generated using the maximin strategy. A suboptimized variant, *HPT-GFP_Sub*, which exhibited a lower predicted TPM than the original sequence, was also generated by applying a model with shuffled codon weights.

A CDS of *picALuc2.0-E50K* (S36) (hereafter “*picALuc*”) was generated by randomly chosen synonymous codons and used to design species-specific optimized variants (*picALuc_At*, *_Ta*, *_Mm*, *_Sc*, and *_Ec* for *A. thaliana*, *T. aestivum*, *M. musculus*, *S. cerevisiae*, and *E. coli*, respectively) by maximizing their predicted TPM values.

### Plasmid construction and gene synthesis

DNA fragments encoding *picALuc* variants (_At, _Ta, _Mm, _Sc, and _Ec) and *HPT-GFP* variants (_Sub, _AtAc, and _ScEc) were synthesized and cloned into pESC-TRP and pGEX-5X-1 vectors, respectively, by GenScript (Piscataway, NJ, USA). Sequences of these variants are provided in table S4, and the correspondence between constructs and specific experiments as well as their generation methods is summarized in table S5. Primers used are listed in table S6.

### Transformation and selection in *Escherichia coli*

For constructs based on pGEX-5X-HPT-GFP, competent *E. coli* DH5α cells were prepared as previously described (S37). Transformation was performed using 1 ng of DNA per 10 μL of competent cells. For pET32-HPT-GFP constructs, *E. coli* BL21(DE3) (Champion21, SMOBIO Technology, Taiwan) was used with 50 ng of DNA per 10 μL of cells. Heat shock was performed at 42°C for 50 seconds, followed by recovery in LB medium for 1 h. Half of the cell suspension was plated onto LB agar containing carbenicillin (Carb, 50 mg/L) to select for the plasmid, while the other half was plated onto LB agar containing hygromycin B (Hyg, 300 mg/L). Colonies were counted after incubation at 37°C for 16 h (for DH5α and Carb-selected BL21(DE3)) or 36 h (for Hyg-selected BL21(DE3)). Hygromycin-resistant colonies were obtained for pET32-HPT-GFP_ScEc even in the absence of IPTG, suggesting that leaky expression of HPT-GFP_ScEc was sufficient to confer resistance.

### Observation of GFP signals in *E. coli*

Single colonies were grown overnight in LB medium with Carb (50 mg/L) at 37°C. The OD_600_ of each culture was measured after 10-fold dilution and confirmed to be approximately 3.9-4.2 for all constructs, indicating that the cells had entered stationary phase with no observable growth bias between the constructs. Cells from 1 mL of culture were harvested by centrifugation, and GFP fluorescence was observed on a blue light transilluminator. Images were captured using a smartphone camera equipped with a GFP filter for stereomicroscopy as previously described (S38). Three independent colonies were analyzed as biological replicates for each construct. GFP signals were detectable for pET32-HPT-GFP_ScEc even in the absence of IPTG, suggesting that leaky expression of HPT-GFP_ScEc was sufficient.

### picALuc assay in *E. coli*

pESC-Trp-picALuc constructs were transformed into BL21(DE3) and resulting single colonies were grown overnight and confirmed to have entered stationary phase (with no growth differences among the constructs) as described above. The culture was then diluted with LB medium to adjust their OD_600_ to 3.8. A 1-mL aliquot of resulting culture was transferred to a sample tube and harvested by centrifugation at 15,000× *g* for 1 min. After discarding the supernatant, the cell pellets were resuspended in 20 µL of a coelenterazine reaction solution, which was prepared by mixing 99 µL of 1× TBS (150 mM NaCl, 20 mM Tris-HCl, pH 7.6) and 11 µL of 1 mM coelenterazine h (reconstituted from 1 mg of powder in 2.5 mL of 99.5% ethanol) (Fujifilm Wako Pure Chemical Co., Osaka, Japan). (The final concentrations in the reaction mixture were 0.1 mM coelenterazine h, 135 mM NaCl, and 18 mM Tris-HCl (pH 7.6).) Bioluminescence was captured using an ImageQuant LAS 4000 Mini (GE care). While luminescence from the *E. coli*-optimized (“Ec”) construct was detectable within a few seconds, a 1-minute exposure time was applied for all samples for subsequent quantification. Data analysis was performed using ImageJ. For each reaction, the mean signal intensity was obtained from three points selected within the signal-harboring area and three points outside the area to define the sample and background signals, respectively. The final value for each replicate was calculated by subtracting the average background signal from the average sample signal. All experiments were performed using three independent colonies for each construct, representing three biological replicates.

### Yeast transformation and hygromycin resistance assay

The Saccharomyces cerevisiae strain BY4741 was transformed using the lithium acetate method according to the Matchmaker protocol (Takara Bio). Competent cells (50 μL each) were incubated with 500 ng of plasmid DNA, 5 μL of carrier DNA (Takara Bio), and 500 μL of PEG/LiAc solution (40% (w/v) PEG4000, 100 mM lithium acetate, 10 mM Tris-HCl, 1 mM EDTA, pH 8.0) at 28°C for 30 min, followed by the addition of 20 μL of dimethyl sulfoxide (DMSO) and heat shock at 42°C for 20 min. After a 4-h recovery in YPGal medium (20 g/L polypeptone, 10 g/L yeast extract, 20 g/L galactose) at 28°C, transformants were plated onto SGal-Ura agar medium (6.7 g/L Yeast Nitrogen Base without amino acids (Difco; BD Biosciences, Franklin Lakes, NJ), 0.78 g/L DO Supplement -Ura (Takara Bio), 20 g/L galactose, 20 g/L agar) with or without 200 mg/L hygromycin B and incubated at 28°C for 5 days.

### picALuc assay in yeast

The *S. cerevisiae* strain W303-1B (BY4949) was obtained from the National BioResource Project (NBRP) Yeast and transformed with pESC-Trp-picALuc constructs as described above. The transformed cells were selected on SGal-Trp agar medium (6.7 g/L Yeast Nitrogen Base without amino acids (Difco), 0.74 g/L DO Supplement -Trp (Takara Bio), 20 g/L galactose, 20 g/L agar). For each construct, three independent colonies of the transformants were inoculated into SGal-Trp liquid medium and cultured for 2 days at 28°C. The cultures (initially around OD_600_ 3.0 for all constructs) were diluted with SGal-Trp to an OD_600_ of 0.5 and cultured for 2 h at 28°C. A 3-mL aliquot of each culture was harvested by centrifugation at 10,000× *g* for 5 s, and the supernatant was discarded. The resulting cell pellets were resuspended in 20 µL of a coelenterazine reaction solution (0.1 mM coelenterazine h, 135 mM NaCl, and 18 mM Tris-HCl (pH 7.6)). Chemiluminescence was captured using an ImageQuant LAS 4000 Mini. A 1-minute exposure time was applied for all samples used for quantification. Data analysis was performed as described above for *E. coli*. Under these conditions, all variants exhibited relatively weak bioluminescence with no significant differences among them.

### GFP and picALuc assays using onion and bok choy

Transient expression in onion (*Allium cepa*) scales was performed using the TSGMAC method (S36). Gold particles (0.6 μm, 60 mg/mL, 7.5 μL) were coated with 250 ng of DNA with a codon variant (pER8d-35S-HPT-GFP or pER8d-35S-GFP) and 250 ng of pBS-35SMCS-mCherry (S39) by adding an equal volume of 50% (w/v) PEG3350 containing 20 mM MgCl_2_. After 5 min of incubation, particles were washed with 70% and 99.5% ethanol, resuspended in 16.5 μL of 99.5% ethanol, and dispersed by sonication. Five microliters of the suspension was bombarded three times into onion scales using TSGMAC. After 16 h of incubation at 28°C, GFP signals were observed using an Olympus BX51 upright fluorescence microscope (S38). For quantification, fluorescence images of GFP and mCherry were captured using a 10× objective with a narrow-band filter (1 s exposure, with no ND filter for GFP; with a U-25ND25 filter for mCherry). The mean gray value was measured for three arbitrary cytoplasmic regions in cells expressing both signals using ImageJ. The average GFP/mCherry signal ratio was calculated to evaluate the relative expression strength among codon variants. The same analysis was performed with bok choy (*Brassica rapa* subsp. *chinensis*), except that a “guide barrel” (S40) was used and images for both signals were captured using the U-25ND25 filter.

A dual-luciferase assay using picALuc variants was performed in bok choy via particle bombardment as previously described (S40) with modifications. To normalize for transfection efficiency, 250 ng of pBS-35S-FLuc (S40) and 400 ng of a *picALuc* variant construct were co-transformed. After incubating a bombarded leaf for 16 h at room temperature, FLuc activity was measured. To quench the FLuc signals, the sections were incubated in 100 mM EDTA (pH 8.0) with 1% (v/v) Triton X-100 for 30 min at room temperature. picALuc signals were then detected using a coelenterazine reaction solution (0.1 mM coelenterazine h, 135 mM NaCl, 18 mM Tris-HCl (pH 7.6), and 1 mM DTT). The picALuc signal intensity was divided by the FLuc signal intensity for each sample. Relative activity was then calculated as the fold-change compared to the mouse-optimized variant (“Mm”), which was set to 1.0.

### Quantitative reverse transcription-PCR (qRT-PCR)

Bok choy leaf cells were co-transformed with 250 ng of pMKV060 (Addgene plasmid #133315; a gift from Daniel Voytas) (S41) as an internal control, and either 250 ng of a *GFP* variant or 400 ng of a *picALuc* variant construct. After incubation for 16 h at room temperature, a leaf section (approximately 1.5 × 1.5 cm) was excised and stored at -80°C. Total RNA was extracted using the NucleoSpin RNA Plant Mini kit (Macherey-Nagel, Düren, Germany), and cDNA was synthesized from 1 μg of total RNA using PrimeScript IV 1st strand cDNA Synthesis Mix (Takara Bio). The resulting cDNA was diluted 10-fold and used as a template for qRT-PCR. Reactions (20 μL) were performed using the StepOne Real-Time PCR System (Thermo Fisher Scientific) and TB Green Premix Ex Taq II FAST qPCR (Takara Bio) with the primers listed in Table S5. To quantify all variants uniformly, a portion of the *NOS* terminator was targeted for amplification. Relative mRNA expression levels were calculated using the comparative cycle threshold method, with *FLuc* serving as the internal reference gene. Each biological replicate consisted of a single bok choy leaf, where all tested variant constructs (three for *GFP* and five for *picALuc*) were bombarded into different areas of the same leaf. The entire experiment was performed in three independent biological replicates.

### picALuc assay using in vitro transcription and translation (IVTT) systems

In vitro protein synthesis was performed using the TNT Coupled Reticulocyte Lysate (RL) and TNT Coupled Wheat Germ Extract (WGE) Systems (Promega, Madison, WI, USA) according to the manufacturer’s instructions, with or without the addition of Recombinant RNase Inhibitor (15 U/reaction; Takara Bio). Each 12-μL reaction contained 10 μL of the TNT premix (including RL or WGE, amino acid mixtures and T7 RNA polymerase) and 100 ng of pESC-Trp-picALuc construct. The reaction mixtures were incubated at 28°C for 1 h. For bioluminescence detection, 2 μL of each IVTT product was mixed with 8 μL of a coelenterazine reaction solution (0.1 mM coelenterazine h, 135 mM NaCl, 18 mM Tris-HCl (pH 7.6); final coelenterazine h concentration in the resulting solution: 80 μM). Chemiluminescence signals were captured using an ImageQuant LAS 4000 Mini (GE care) with exposure times of 3-5 min. Signal intensities were quantified using ImageJ by averaging the mean gray values of three arbitrary points within each reaction, after subtracting the background signal. Each assay was performed in three independent replicates.

### Statistical analysis of experimental data

Statistical analyses and data visualization were performed using Python (v3.12) with the scipy, statsmodels, pandas, and seaborn libraries. To evaluate differences among multiple codon variants, the non-parametric Kruskal-Wallis test was employed. When a significant difference was detected (*P* < 0.05), Tukey’s Honestly Significant Difference (HSD) test was performed as a post-hoc analysis for pairwise multiple comparisons. All data are presented as the mean ± SD, with individual data points displayed as a strip plot. For qRT-PCR analysis, statistical tests were performed on log_10_-transformed values. A *P*-value of less than 0.05 was considered statistically significant.

---

**Figure S1.**
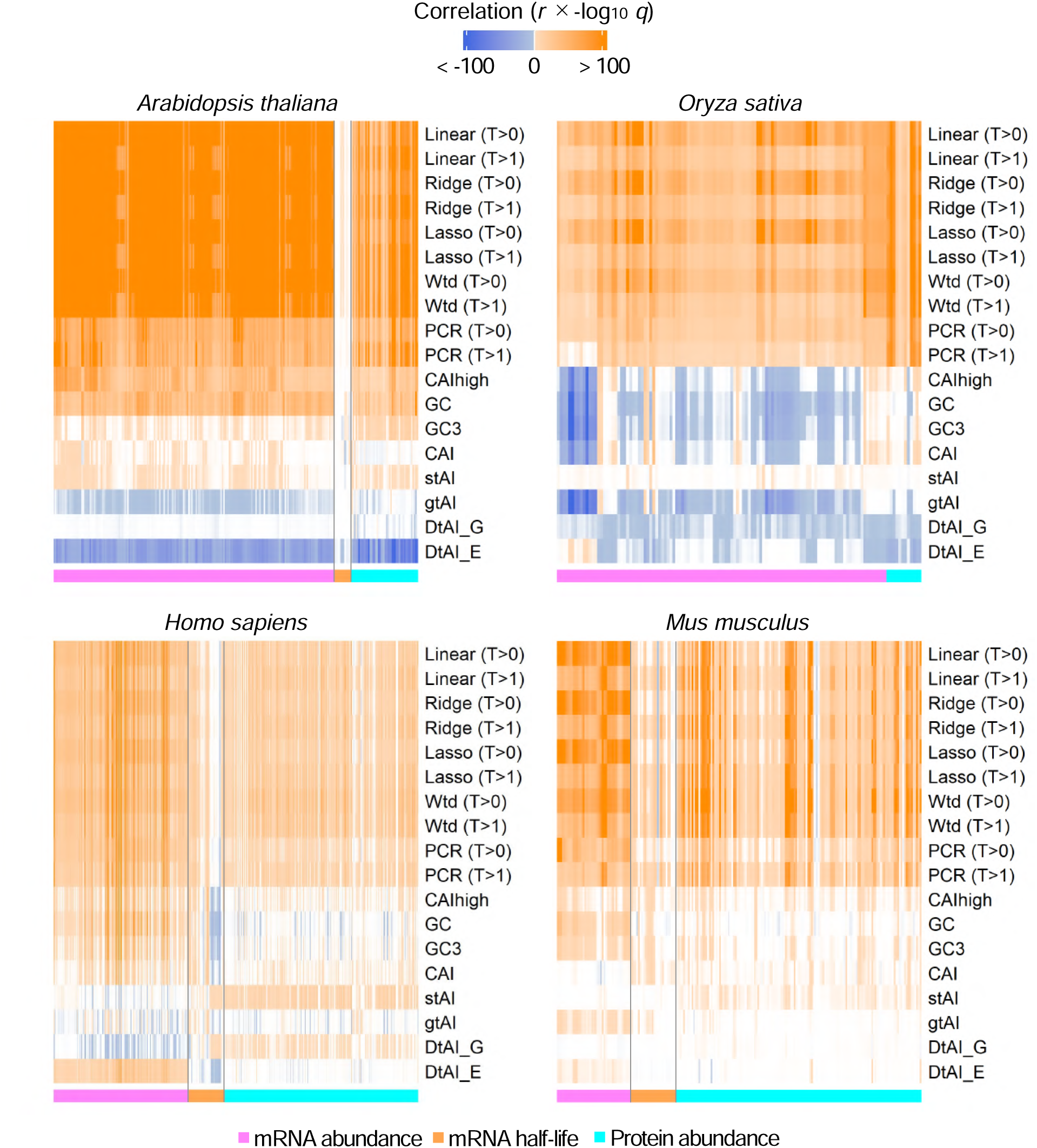
Comparison of regression strategies for predicting mRNA abundance and physiological features from codon usage. Values of Pearson’s *r* multiplied by -log_10_ *q* (FDR-corrected *P* values) were calculated across different regression models and codon-related indices. Models were trained using mean log1p-transformed TPM as the response variable and centered log-ratio (CLR)-transformed codon frequencies as explanatory variables. Five regression methods were evaluated: simple linear (Linear), Ridge, Lasso, weighted (Wtd.), and principal component (PCR). The predictive performance of these models, alongside established indices (i.e., GC and GC3 contents and tAI- and CAI-based metrics), was assessed by correlating their outputs with observed log_10_(TPM), mRNA half-life, log_10_(protein abundance) from PaxDb. T>0 and T>1 denote gene filtering thresholds based on TPM levels. CAIhigh represents the index calculated from the top 5% highly expressed genes. Species names are indicated at the top of the panels.

**Figure S2.**
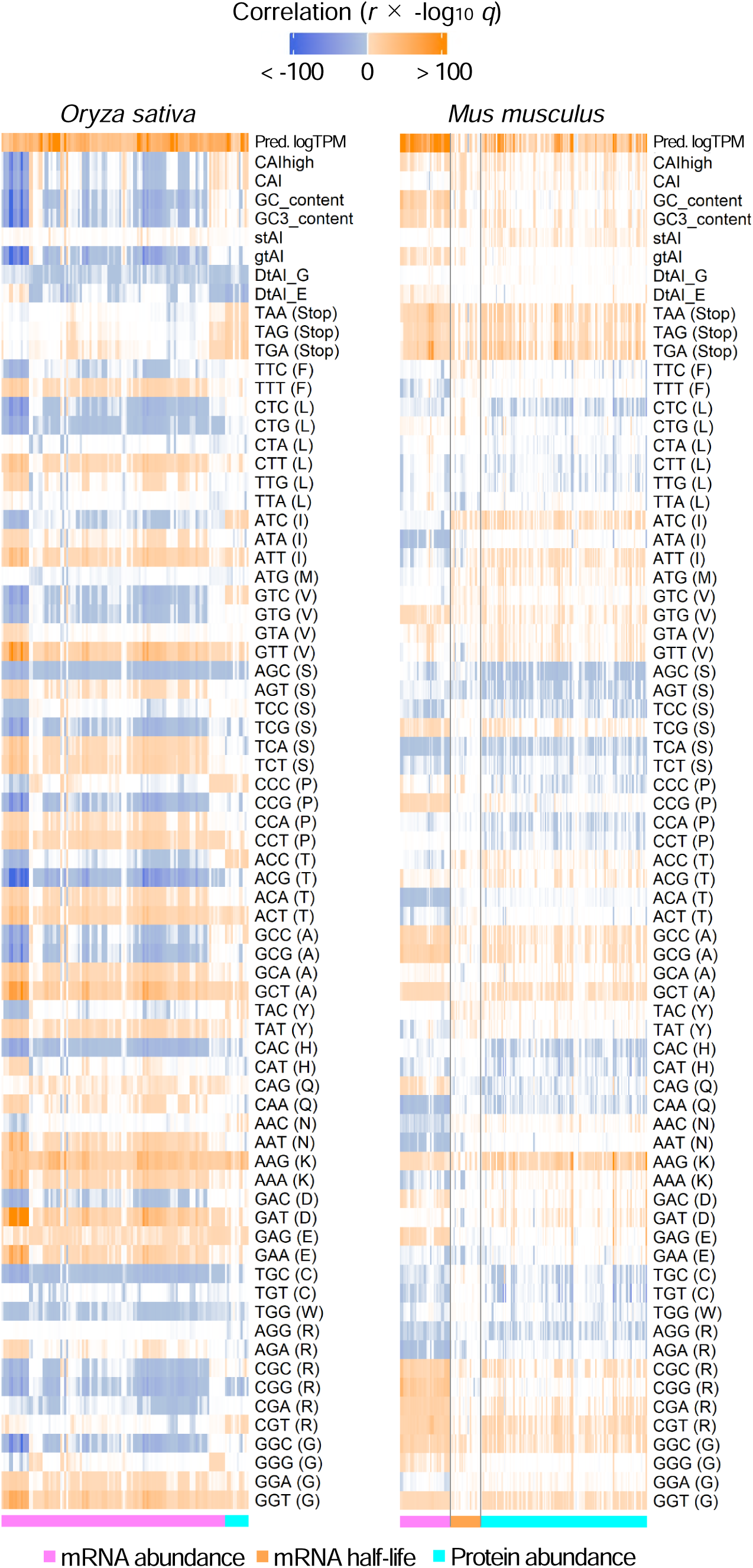
Correlations between codon-related metrics and expression levels in rice and mouse. Values of Pearson’s *r* multiplied by -log_10_ *q* (FDR-corrected *P* values) were calculated from correlations of predicted log_10_(TPM) (Pred. logTPM), codon-related indices, and individual codon frequencies with observed log_10_(TPM), mRNA half-life (for mouse), and log_10_(protein abundance). Predicted values were obtained from simple linear regression models trained on genes with TPM > 0. Species names are indicated at the top of the panels

**Figure S3.**
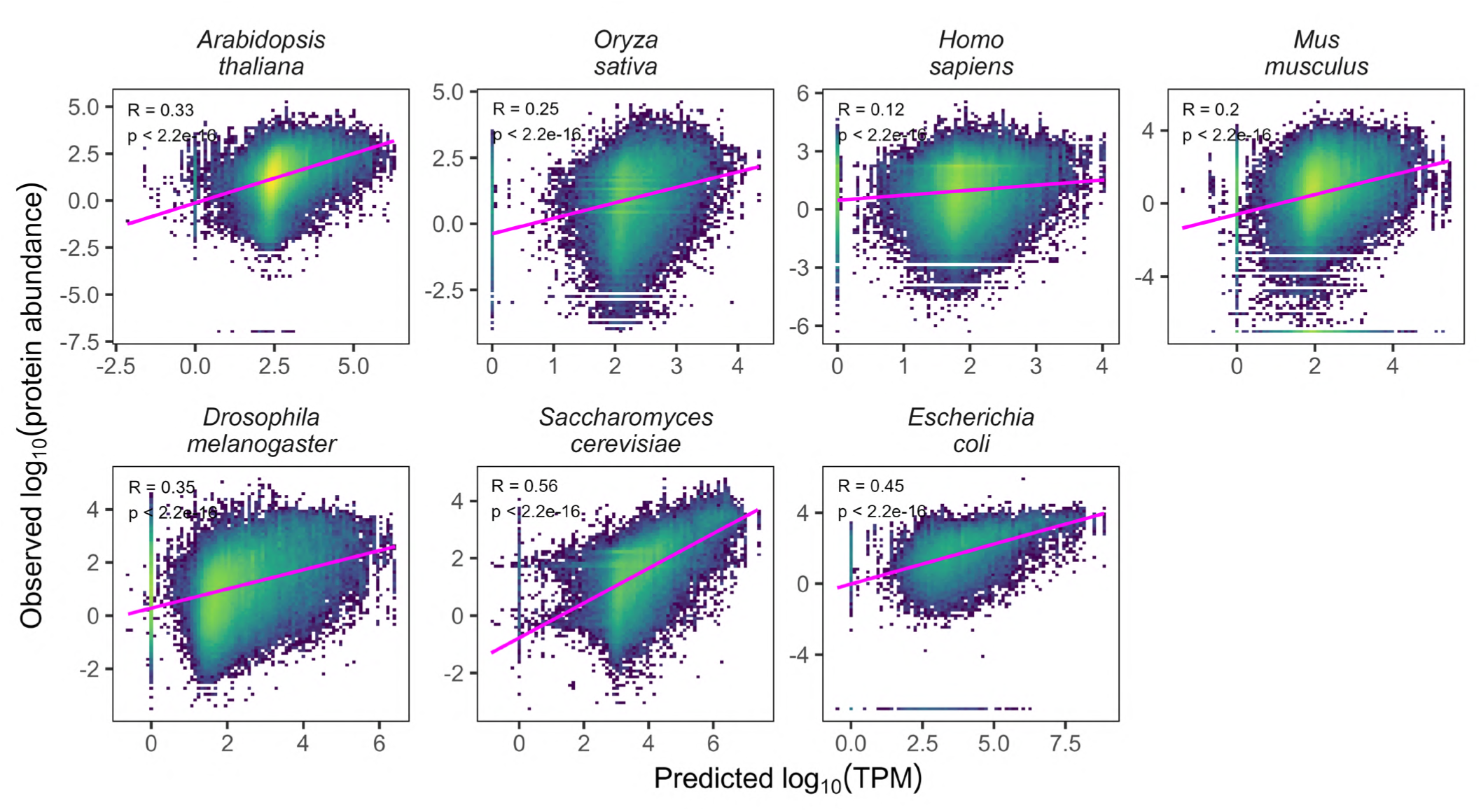
Correlations between predicted log_10_(TPM) and log_10_(protein abundance). Predicted log_10_(TPM) were derived from simple linear regression models based on codon frequencies. Observed protein abundance data were obtained from PaxDb. Species names are indicated at the top of the panels. Pearson’s correlation coefficient (R) and *P* values (p) are indicated within each panel.

**Figure S4.**
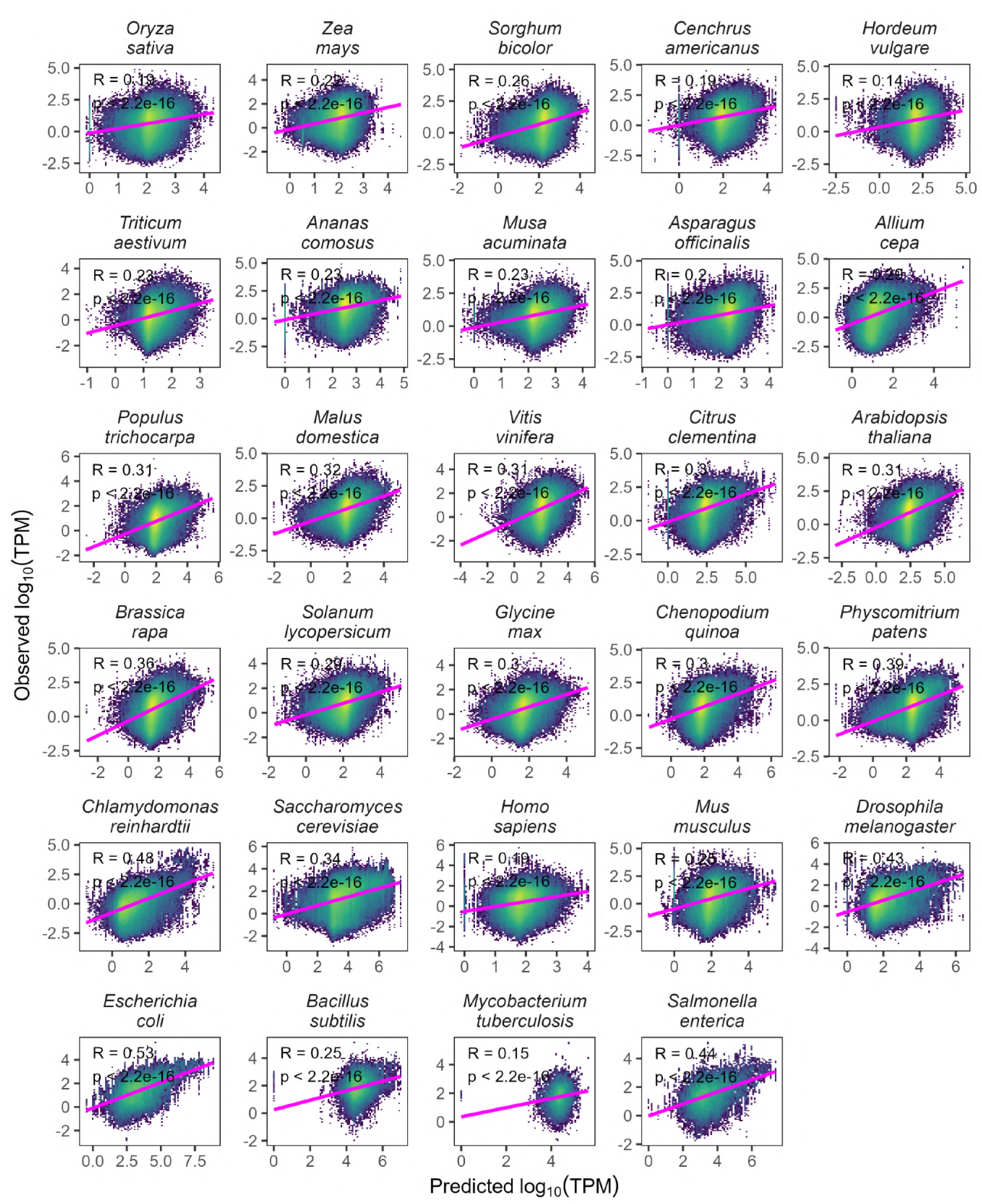
Correlations between predicted log_10_(TPM) and observed log_10_(TPM) across 29 species. Predicted log_10_(TPM) values were derived from simple linear regression models based on codon frequencies. Species names are indicated at the top of the panels. Pearson’s correlation coefficient (R) and *P* values (p) are indicated within each panel.

**Figure S5.**
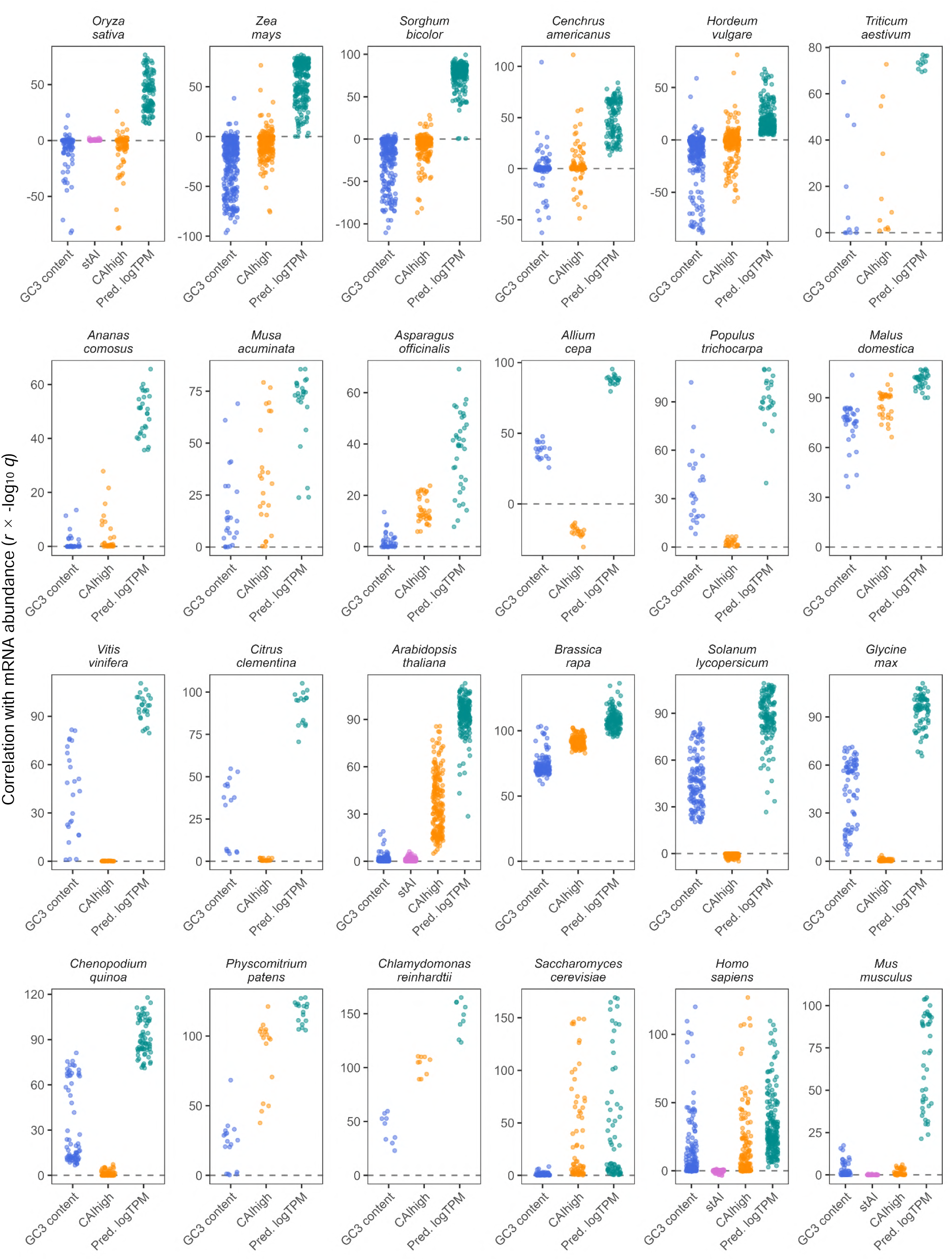
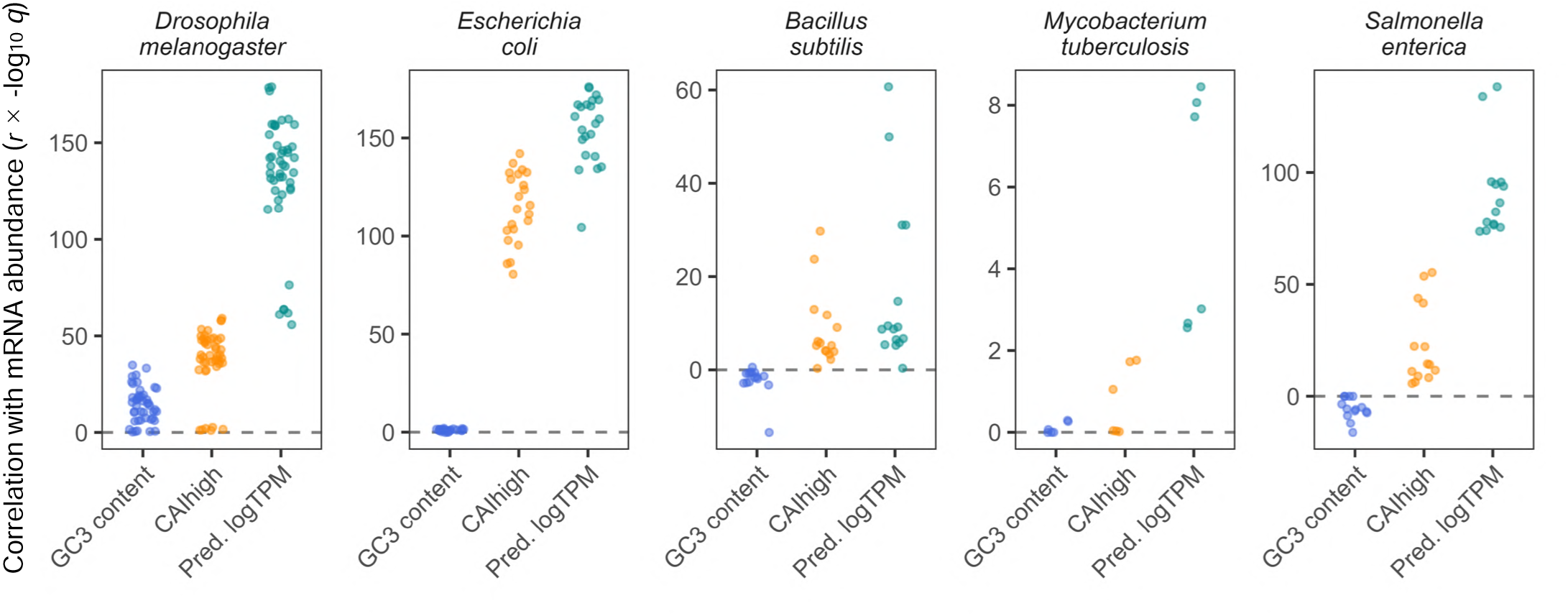
Correlations between codon-based indices and observed log_10_(TPM) across 29 species. Values of Pearson’s *r* multiplied by -log_10_ *q* (FDR-corrected *P* values) were calculated from correlations of GC3 content, stAI (only for *A. thaliana*, *O. sativa*, *H. sapiens*, and *M. musculus*), CAIhigh, and predicted log_10_(TPM) (Pred. logTPM) with observed log_10_(TPM). Observed log_10_(TPM) was used as the reference for all correlations. Species names are indicated at the top of the panels.

**Figure S6.**
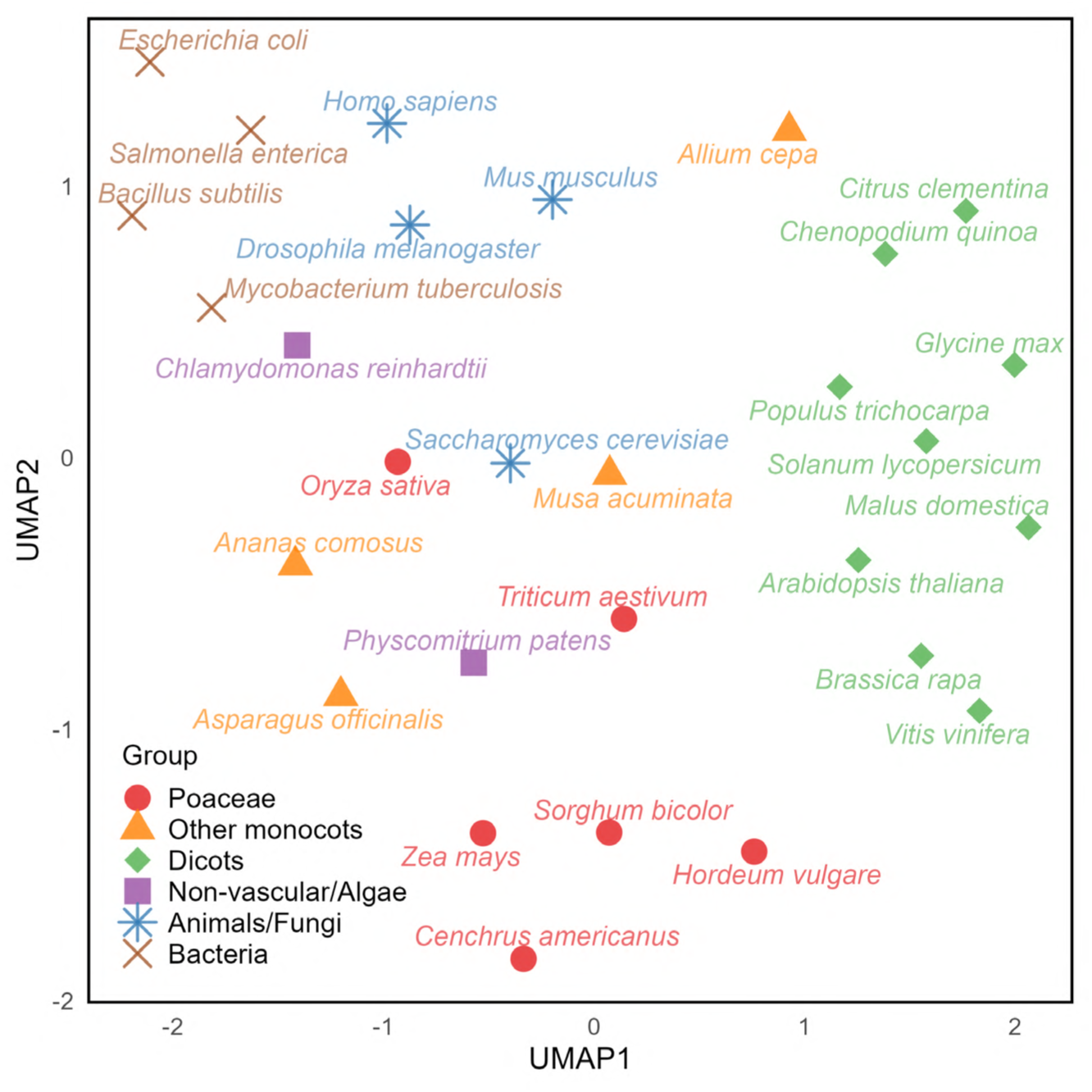
UMAP of species based on codon weights. Each point represents an individual species, and their positions were determined using the coefficients obtained from simple linear regression models. Proximity between points indicates similarity in codon weight profiles across the 29 species.

**Figure S7.**
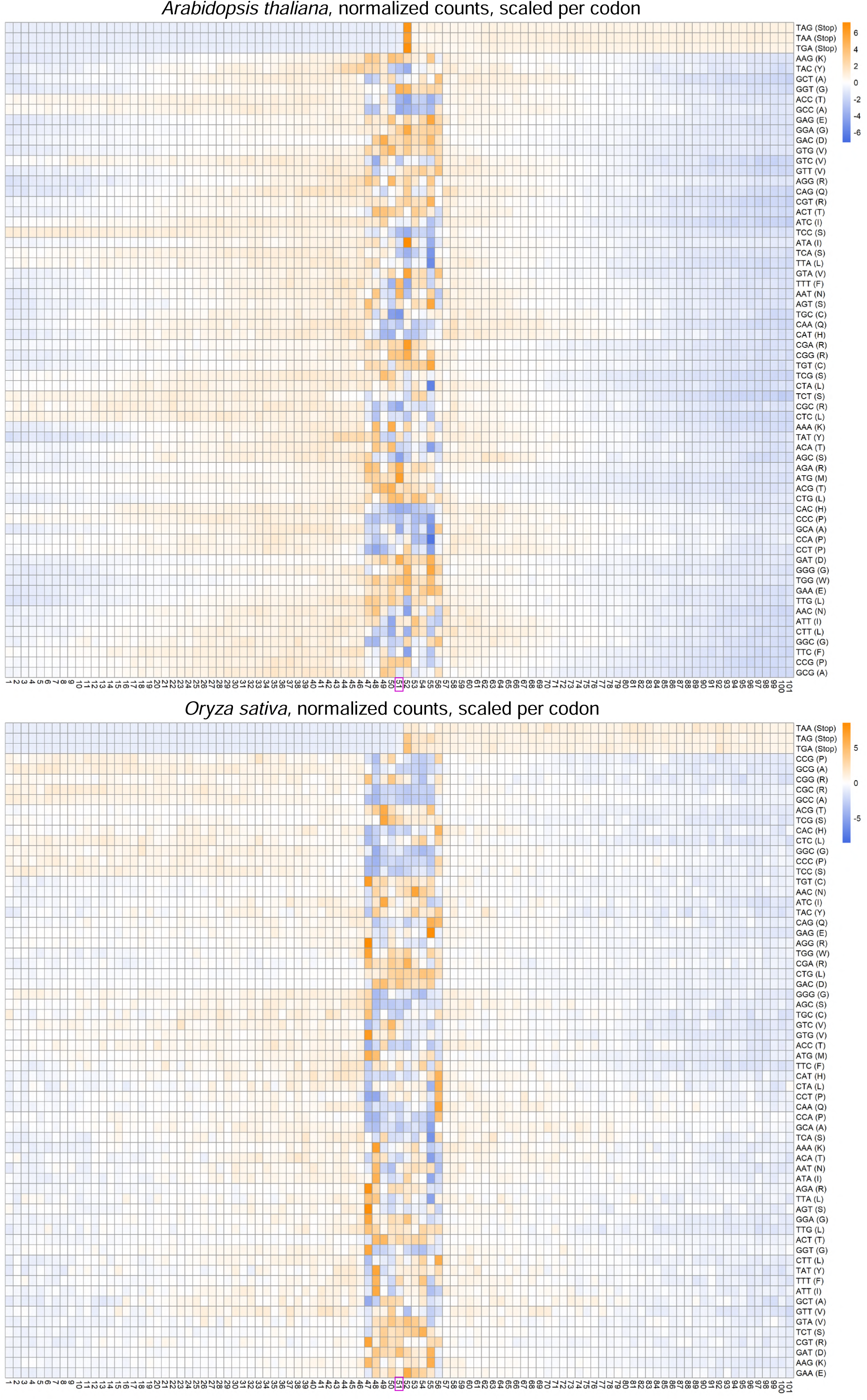
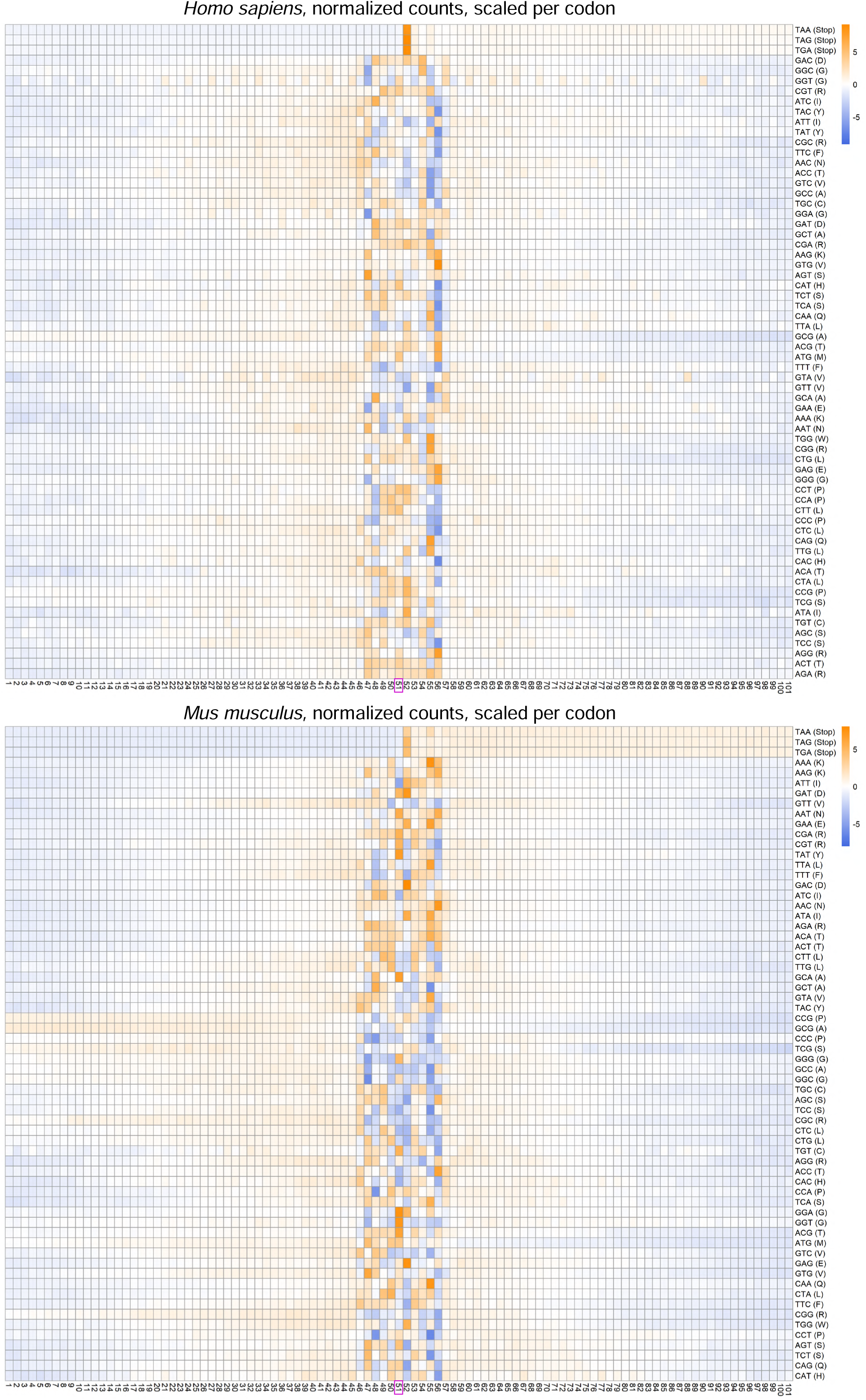
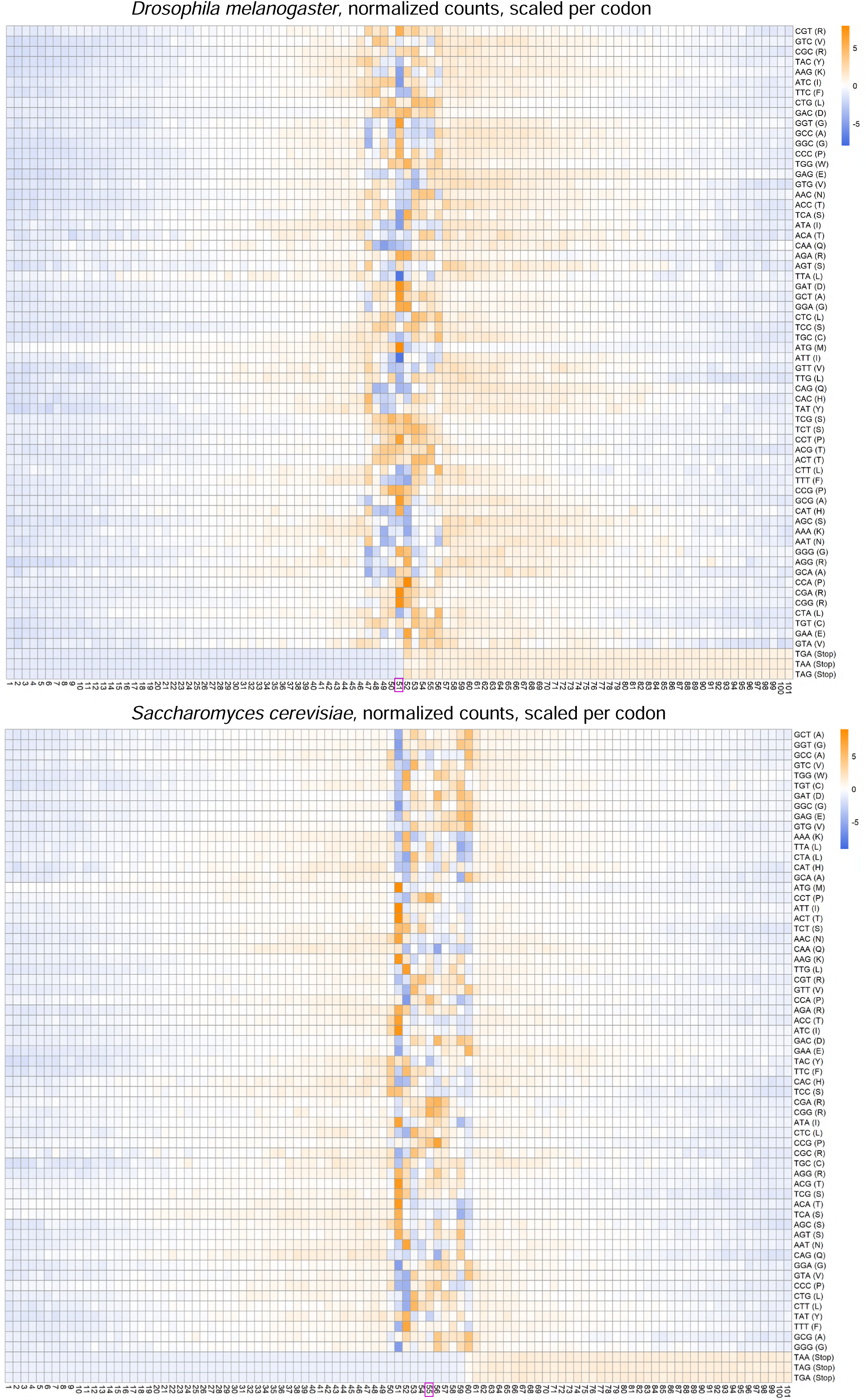
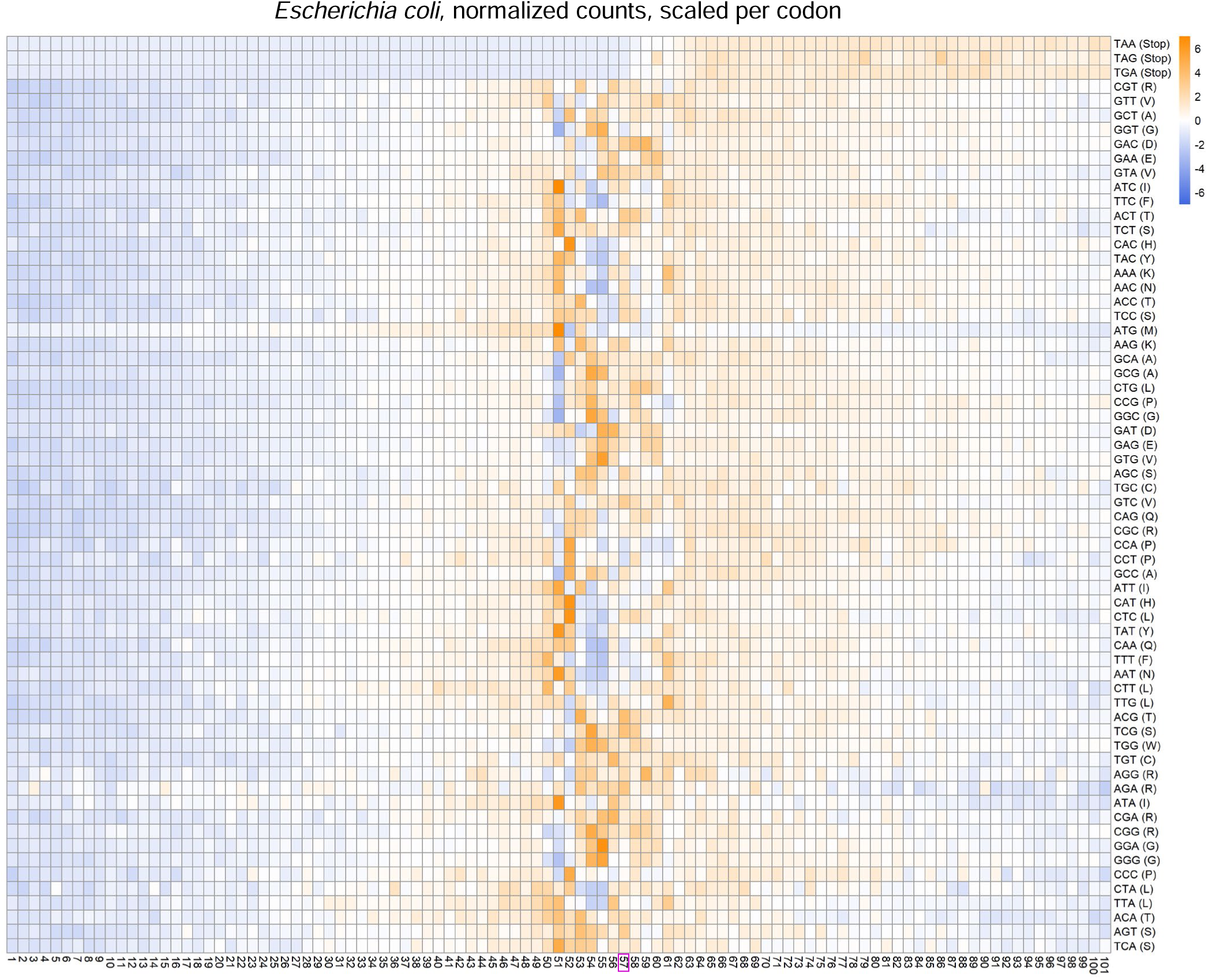
Ribo-seq-weighted cumulative codon frequency patterns across seven species. Ribo-seq-weighted cumulative codon frequencies were calculated across a 101-codon window centered on the designated reference sites (offsets) for Ribo-seq reads. At each position within a 101-codon window centered on reference sites (offsets), P-site read counts were normalized by the mean read density of each gene, summed up for each codon across all genes, and subsequently divided by the total occurrence of each codon across all analyzed genes. The resulting values were scaled by codon and subjected to clustering to generate the heatmaps. For *S. cerevisiae* and *E. coli*, the reference sites were adjusted based on available CDS coordinates due to the lack of sufficient 5’ and 3’ untranslated sequence data, resulting in a shift in the putative P-site (boxed at the bottom) compared to other species. Species names are indicated at the top of the panels.

**Figure S8.**
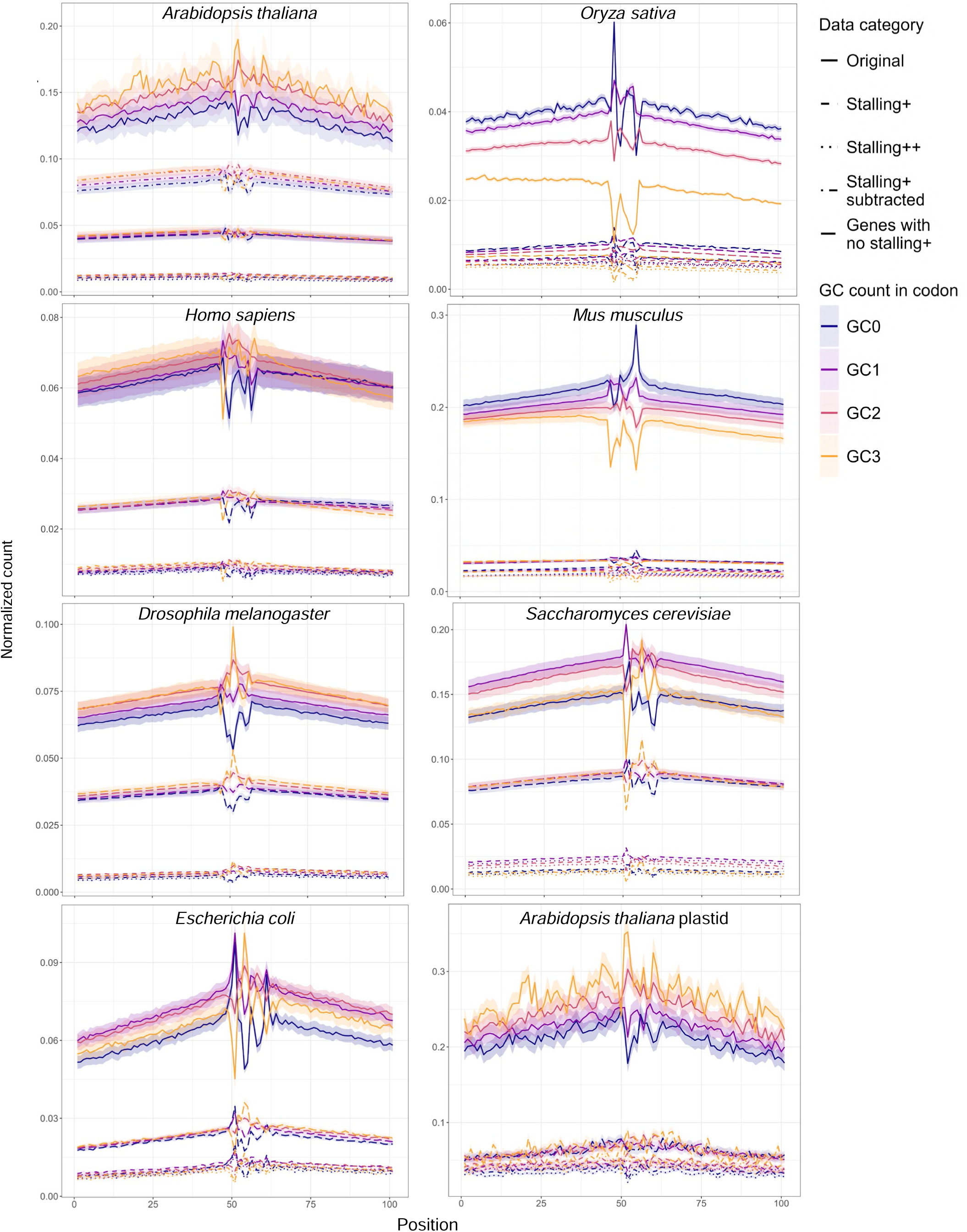
Ribo-seq-based positional frequency patterns of GC content-based codon groups across seven species. Ribo-seq-weighted cumulative codon frequencies were aggregated for codon groups with different GC contents (0, 1, 2, or 3 GC bases) across a 101-codon window. These values were calculated independently for each Ribo-seq sample and are presented as ribbon plots representing the range between the minimum and maximum values, with the middle line indicating the mean across samples. Sites with significantly high read counts were identified using two significance thresholds and designated as stalling+ and stalling++. Ribbons are also provided for those data, stalling+ subtracted data (where specific stalling+ sites were removed) and genes with no stalling+ (where entire genes containing at least one stalling+ site were excluded). Species names are indicated at the top of the panels.

**Figure S9.**
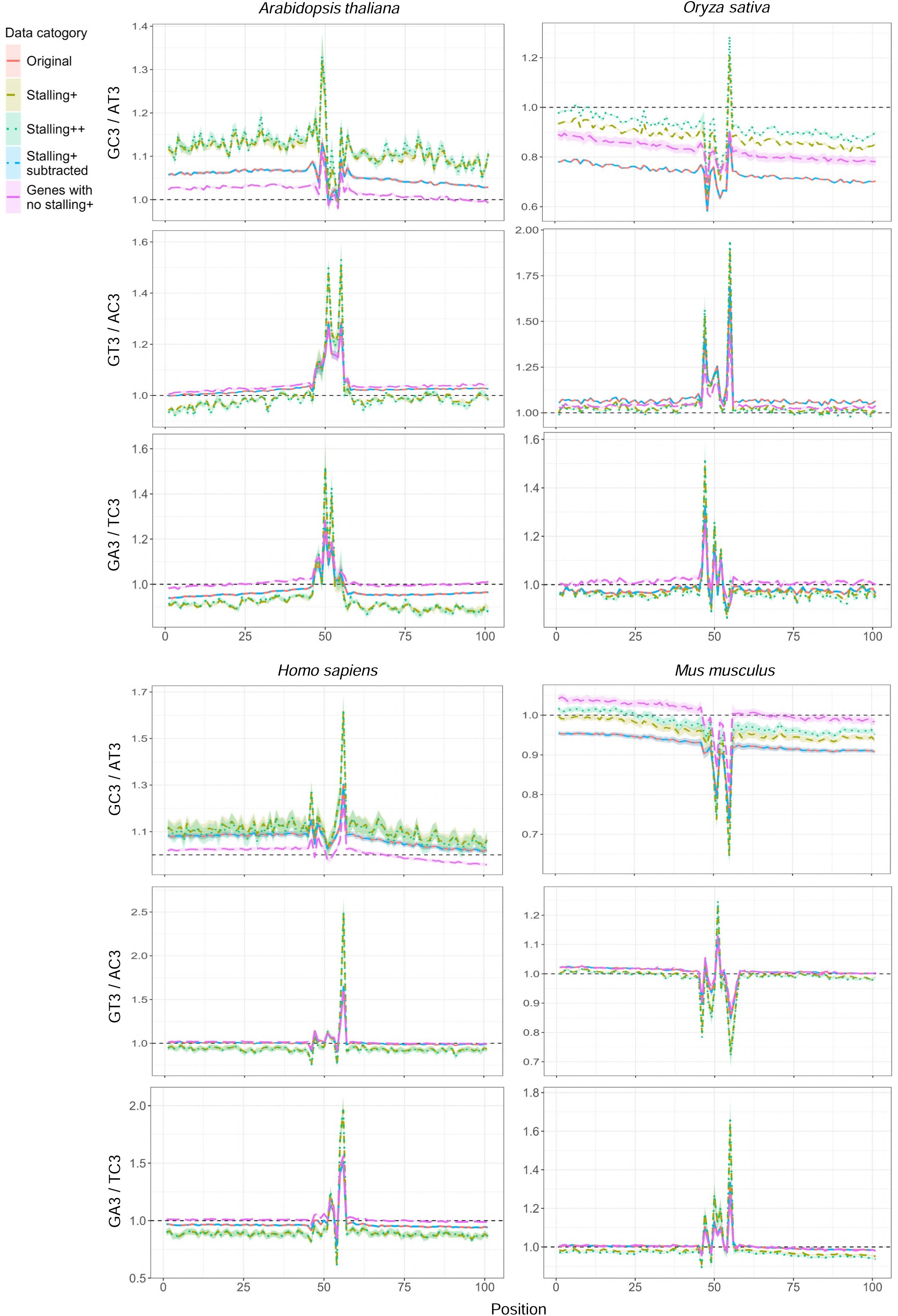
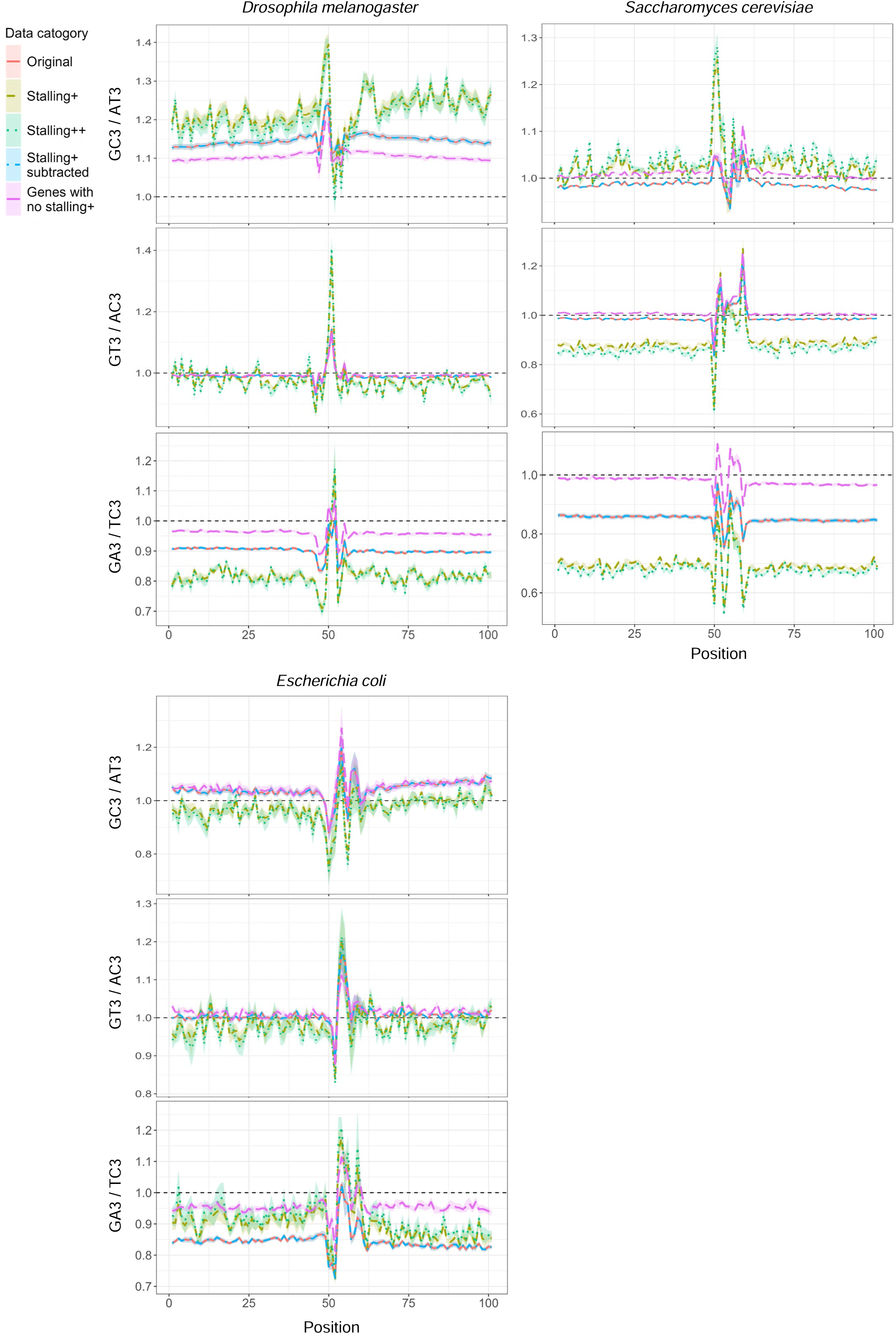
Codon nucleotide composition ratios based on Ribo-seq. Values of nucleotide composition ratios at the third codon position, (GC3 / AT3), (GT3 / AC3), and (GA3 / TC3) were calculated across a 101-codon window using Ribo-seq-weighted cumulative codon frequencies. These ratios were determined independently for each Ribo-seq sample and are presented as ribbon plots representing the range between the minimum and maximum values, with the middle line indicating the mean across samples. Sites with significantly high Ribo-seq read counts were identified using two significance thresholds and designated as stalling+ and stalling++. Ribbons are also provided for those data, stalling+ subtracted data (where specific stalling+ sites were removed) and genes with no stalling+ (where entire genes containing at least one stalling+ site were excluded). Species names are indicated at the top of the panels.

**Figure S10.**
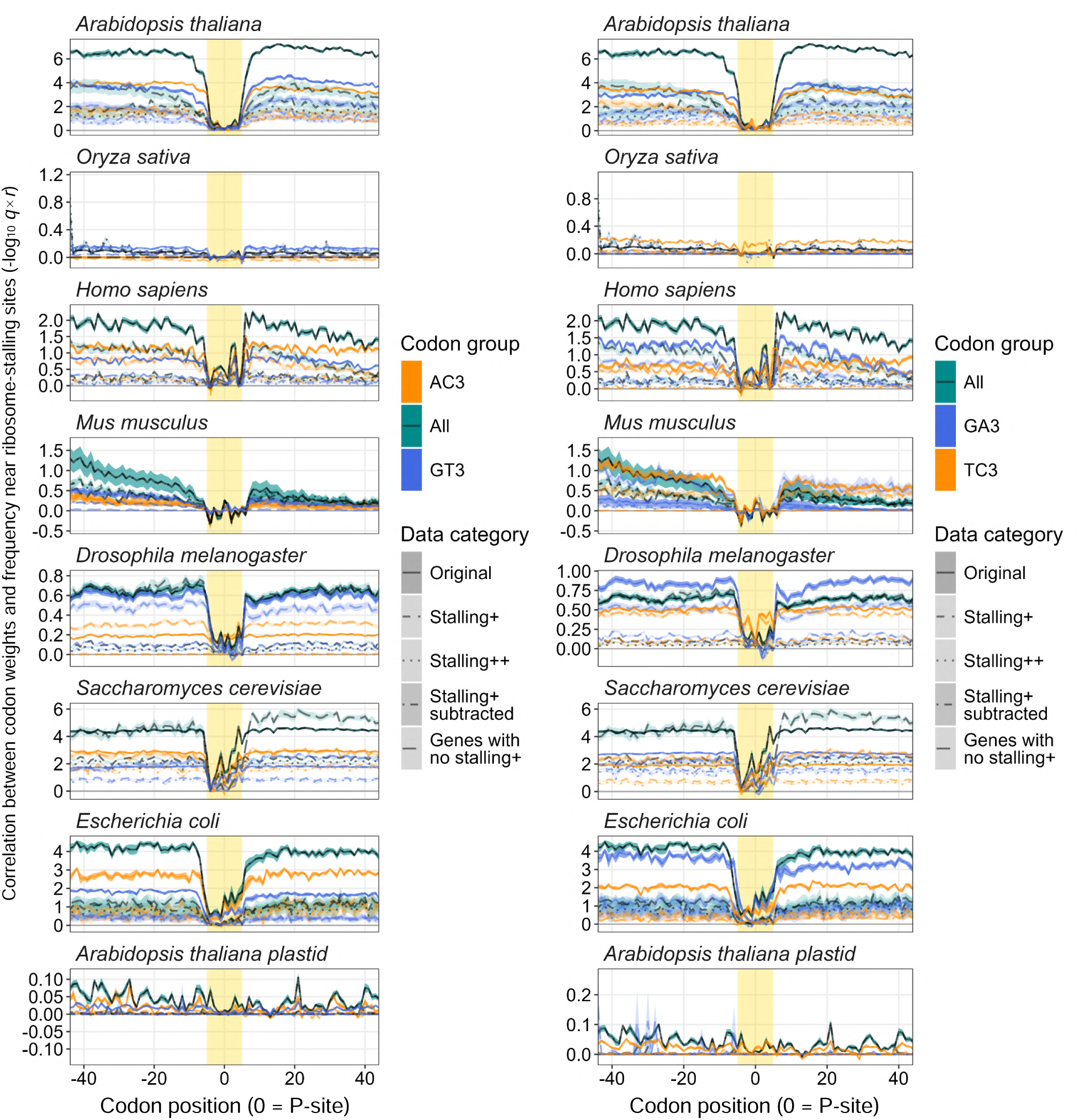
Positional correlations between codon weights and Ribo-seq-weighted codon frequencies. Values of Pearson’s *r* multiplied by -log_10_ *q* (FDR-corrected *P* values) were calculated by correlating the codon weights (from simple linear regression models trained on genes with TPM > 0) with the Ribo-seq-weighted cumulative codon frequencies across an 81-codon window. For each position, the correlation was tested using the 61 non-stop codons. These values were determined independently for each Ribo-seq sample and are presented as ribbon plots representing the range between the minimum and maximum values, with the middle line indicating the mean. The putative ribosome-occupied regions were estimated by the codon frequency patterns and the putative P-sites presented in fig. S7 and are highlighted by background color. Species names are indicated at the top of the panels. Similar plots for the AT3 and GC3 codon groups are provided in fig. 4A.

**Figure S11.**
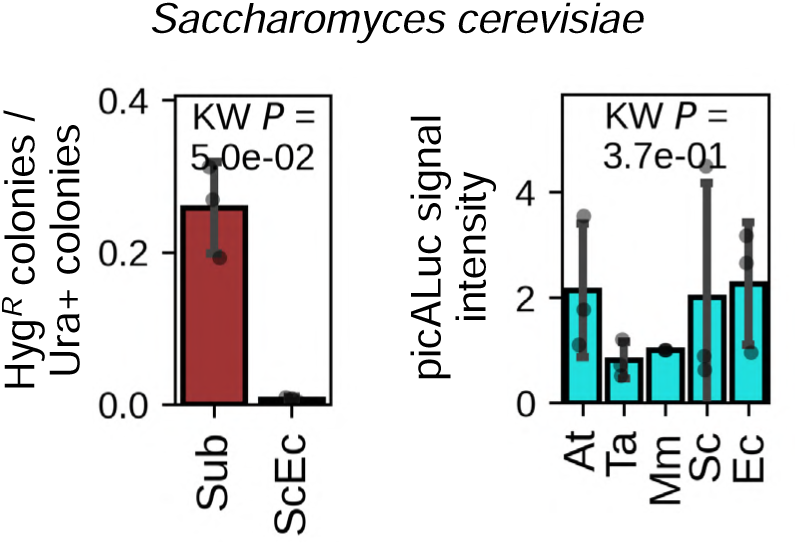
Functionality of *HPT-GFP* and *picALuc* codon variants in *Saccharomyces cerevisiae*. (Left) Two *HPT-GFP* codon variants, Sub (suboptimal) and ScEc (optimized for *S. cerevisiae* and *Escherichia coli*) were transformed into *S. cerevisiae* strain BY4741. Ratios of hygromycin-resistant (Hyg*^R^*) colonies (dependent on *HPT* functionality) and uracil-prototrophic (Ura^+^) colonies (control) are presented. (Right) Five *picALuc2.0-E50K* codon variants, At (optimized for *A. thaliana*), Ta (for *T. aestivum*), Mm (for *M. musculus*), Sc (for *S. cerevisiae*) and Ec (for *E. coli*) were transformed into *S. cerevisiae* strain W303-1B. picALuc bioluminescence signals in cultured transformed cells are presented. For both panels, bars and error bars represent means and SD, respectively, from three biological replicates (indicated by individual dots). KW *P* indicates the *P* value obtained from the Kruskal-Wallis test.

**Figure S12.**
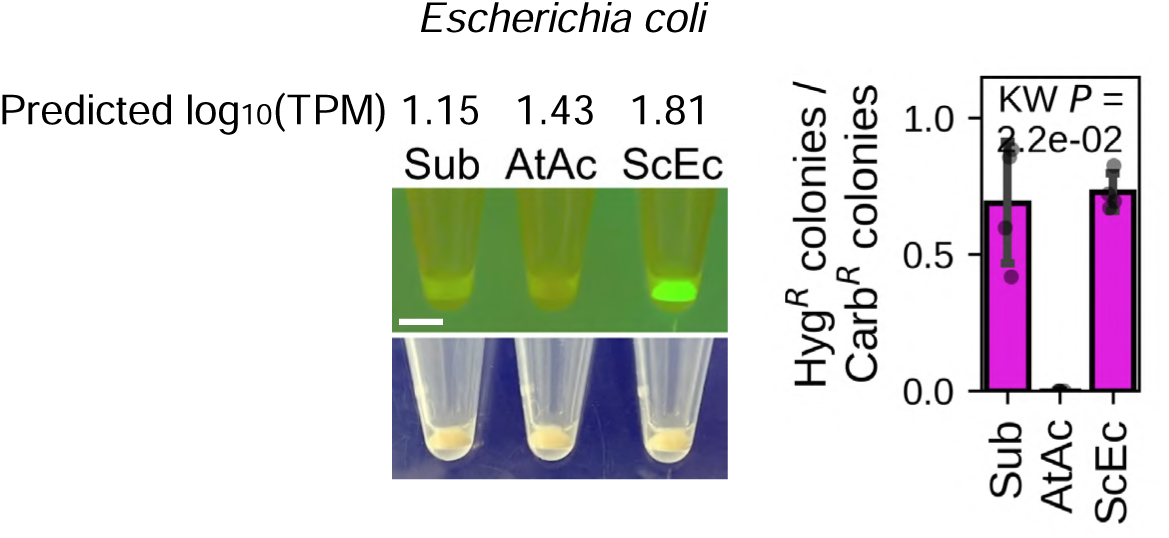
Expression and functionality of *HPT-GFP* codon variants fused to a common *GST* sequence in *Escherichia coli*. Three *HPT-GFP* codon variants, Sub (suboptimal), AtAc (optimized for *A. thaliana* and *Allium cepa*), and ScEc (for *S. cerevisiae* and *E. coli*) with a common GST CDS at their 5’ ends were transformed into *E. coli* strain DH5α. (Left) Predicted log_10_(TPM) values for the GST-HPT-GFP variants, followed by representative GFP fluorescence and bright-field images of transformed cells cultured overnight. Scale bar = 4 mm. (Right) Ratios of hygromycin-resistant (Hyg*^R^*) colonies (dependent on *HPT* functionality) and carbenicillin-resistant (Carb*^R^*) colonies (control) obtained after transformation. Bars and error bars represent means and SD, respectively, from four biological replicates (indicated by individual dots). KW *P* indicates the *P* value obtained from the Kruskal-Wallis test.

## Legends for table S1 to S6

Table S1. Overview of data and analysis summaries.

Table S2. Custom scripts and programs for data analysis and visualization.

Table S3. Curated datasets for tRNA gene numbers, tRNA expression, and mRNA half-lives.

Table S4. Sequences of codon variants used in this study.

Table S5. Constructs and experiments using codon variants.

Table S6. Primers used for construct generation and mRNA qRT-PCR analysis of mRNA abundance.

